# M2WISH: an easy and efficient protocol for whole-mount mRNA *in situ* hybridization that allows 3D cell resolution of gene expression in *Arabidopsis thaliana*

**DOI:** 10.1101/2024.01.18.576197

**Authors:** Liudmila Chelysheva, Halima Morin, Eric Biot, Antoine Nicolas, Philippe Rech, Marco da Costa, Lisa Barel, Patrick Laufs, Jean-Christophe Palauqui

**Author notes:** **Correspondence:** Liudmila Chelysheva, Université Paris-Saclay, INRAE, AgroParisTech, Institut Jean-Pierre Bourgin (IJPB), 78000, Versailles, France. Jean-Christophe Palauqui, Université Paris-Saclay, INRAE, AgroParisTech, Institut Jean-Pierre Bourgin (IJPB), 78000, Versailles, France. **Email adresses:** Liudmila, Halima, Eric, Antoine, Philippe, Marco da, Lisa, Patrick, Jean-Christophe.

## Abstract

Gene expression analysis is essential for understanding the mechanisms involved in plant development. Here, we developed M2WISH, a protocol based on <u>M</u>icro<u>W</u>ave treatment for <u>W</u>holemount mRNA In <u>S</u>itu <u>H</u>ybridization in Arabidopsis. By permeabilizing tissues without damaging cellular organisation this protocol results in high and homogeneous hybridization yields that enables systematic analysis of gene expression dynamics. Moreover, when combined with cellular histochemical staining, M2WISH provides a cellular resolution of gene expression on roots, aerial meristems, leaves and embryos in the seed. We applied M2WISH to study the spatial dynamics of *WUSCHEL* (*WUS*) and *CLAVATA3* (*CLV3*) expression during *in vitro* meristematic conversion of roots into shoot apical meristems. Thus, we showed that shoot apical meristems could arise from two different types of root structures that differed by their *CLV3* gene expression patterns. We constructed 3D cellular representations of *WUS* and *CLV3* gene co-expression pattern, and stressed the variability inherent to meristem conversion. Thus, this protocol generates a large amount of data on the localization of gene expression, which can be used to model complex systems.

## Background

Modern integrative biology studies require fast and efficient tools to visualise the expression of genes in their spatio-temporal context. One method of choice to study the dynamics of gene expression in time and space is to use transgenic reporter constructs coupled to a fluorescent protein. However, these studies require numerous steps to construct and select reporters and introducing them into different genetic contexts is time consuming. Moreover, expression of transgene can be biased on localization either because promoter sequences are incomplete, or integration in ectopic position in the genome induces expression fluctuations. *In situ mRNA* hybridization based on the complementarity of an antisense ribo-probe with specific cellular RNA populations (Gall, 2016; Johnson *et al*., 1991) was originally the technic to reveal the accumulation of mRNAs transcripts in any tissues and genetic context. This detection was classically carried out on sections with a colorimetric dye and is now widely used, despite a laborious protocol. For instance, medium-throughput *in situ* RNA hybridization dataset for 39 genes of a rapidly growing and complex Arabidopsis embryo was successfully generated and complemented the tissue-specific high-throughput transcriptomics with a higher spatial resolution (Francoz *et al*., 2016). The development of fluorescent dye based on the Tyramide Signal Amplification (TSA) system combining with parietal and nuclear markers followed with confocal imaging provided the possibility to investigate the cellular and subcellular mRNAs localization of meristem and flower development genes on sectioned tissue of *Arabidopsis* (Yang *et al*., 2020). To overcome the time-consuming steps of embedding and sectioning, several protocols of Whole Mount *in Situ* Hybridization (WISH) have been developed and optimized for permeable (root, embryo), or accessible samples (apical and floral meristem) (Hejátko *et al*., 2006; Rozier *et al*., 2014). Here, we propose an easy and robust WISH protocol based on Microwave (MW) treatment that is applicable without adaptation to organs as diverse as embryo, root, leaves, apical and floral meristems of Arabidopsis. Because this treatment leads to detection in all tested tissues, it allows unambiguous identification of cells expressing (and cells not-expressing) the gene of interest, thus greatly limiting the risk of false-negative. We use Alkaline Phosphatase amplification (AP) detection system that combined colorimetric and fluorescence signals for macroscopic and microscopic detection. With this protocol, simultaneous gene expression and fluorescent staining of cell wall and nucleus can be achieved at the whole organ level with a cell resolution after confocal imaging. We extend our protocol to double WISH, combining two different detections, AP and peroxidase (POD), yielding in an unambiguous mRNAs colocalization of two different genes. We tested our protocol on samples generated during direct conversion of lateral root meristems into shoot meristems (Rosspopoff *et al*., 2017). In this system, incipient lateral roots rapidly acquire the shape and the identity of a typical Shoot Apical Meristem (SAM). A major transcriptional reprogramming was documented during the course of meristem conversion by analysis of transcriptomics, reporter lines and colorimetric *in situ* hybridization (Rosspopoff *et al*., 2017). Here, we precisely localized *WUSCHEL* (*WUS*) and *CLAVATA3* (*CLV3)* mRNA by double WISH and used a 3D parametric representation of their expression patterns to characterize in detail their evolution during meristem conversion. Thus, we found that *WUS* and *CLV3* are expressed following two stereotypical patterns. One is the canonical pattern described during rosette axillary meristems establishment (Xin *et al*., 2017) where *WUS* and *CLV3* are initially expressed in a central domain while later *CLV3* expression shifts from the central to the apical domain. In the second one, *CLV3* expression domain is initially located to apical position, when *WUS* occupies a central domain. At later stages, all converted meristems show the patterns expected for SAMs with an apical *CLV3* domain and a lower *WUS* domain. Therefore, our analysis reveals a high flexibility in the process of meristem conversion with different developmental paths leading to identical final organization in established meristems.

## RESULTS

### Microwave treatment improves organ permeability and accessibility

The classical protocol for *in situ* mRNAs detection can be divided in four major steps: sample fixation and permeabilization, complementary riboprobe hybridization to cellular mRNAs, *in situ* ribo-probe detection and signal visualization. The major challenge for establishing a WISH protocol robust towards different tissues and organs is to optimize permeabilization treatments to allow uniform access to the different reagents, mostly the ribo-probe and antibodies, while preserving tissue integrity. Enzymatic cell wall digestion is often used to favor permeability of organs and accessibility to the different reagents. However, this step needs to be adapted for each organ and often weakens samples especially during ribo-probe hybridization leading to a considerable loss of yield. We thus decided to move this step after ribo-probe hybridization. Instead, we opted for a microwave (MW) treatment in a series of ethanol dilutions to permeabilize the sample towards the ribo-probes. MW treatment is known to increase the accessibility of protein to antibodies on chromosome spread (Chelysheva *et al*., 2010). We performed the MW in ethanol series because hot ethanol treatment delipidates cuticle and helps to extend the Pseudo-Schiff Propidium Iodide (PS-PI) staining for aerial tissue of all Arabidopsis organs at all stages of development (Truernit *et al*., 2008). We named this protocol M2WISH for <u>M</u>icro<u>W</u>ave treatment for <u>W</u>holemount mRNA In <u>S</u>itu <u>H</u>ybridization.

Thus, we tested this modified protocol on 14-day–old seedlings of Arabidopsis. At this stage plantlets possess a well-developed root system with numerous lateral root primordia and outgrowing secondary roots of different stages with various permeability. We hybridized MW-treated and non-treated plantlets with *PLETHORA1* (*PLT1*) anti-sense probe, a key player in root meristem development (Du and Scheres, 2017), and found that for samples treated with MW, all roots were labeled with *PLT1* in their root meristem initials as expected (100% of lateral roots for 10 plantlets, Figure 1). In contrast, in the same conditions only a part of non-MW-treated roots bore a signal (30 % of the most outgrown roots for 3 plantlets, Figure S1), suggesting that variable tissue permeability could be a bottleneck for efficient mRNA *in situ* hybridization. As a negative control, we hybridized MW treated plantlets with PLT1 sense probe and found no labelling inside root meristems (Figure S2), but some unspecific staining at the cuticle and in some root cap cells. We further validated our protocol on root and shoot meristems by showing a reproducible and efficient labeling of the quiescent or organizing centers by *WUSCHEL-RELATED HOMEOBOX 5* (*WOX5*) and *WUS* respectively (Figure S3, Figure S4). Thus, MW treatment in a series of ethanol homogenizes the tissue permeability and ensures uniform and comprehensive RNA probe accessibility to cells.

**Figure 1.**
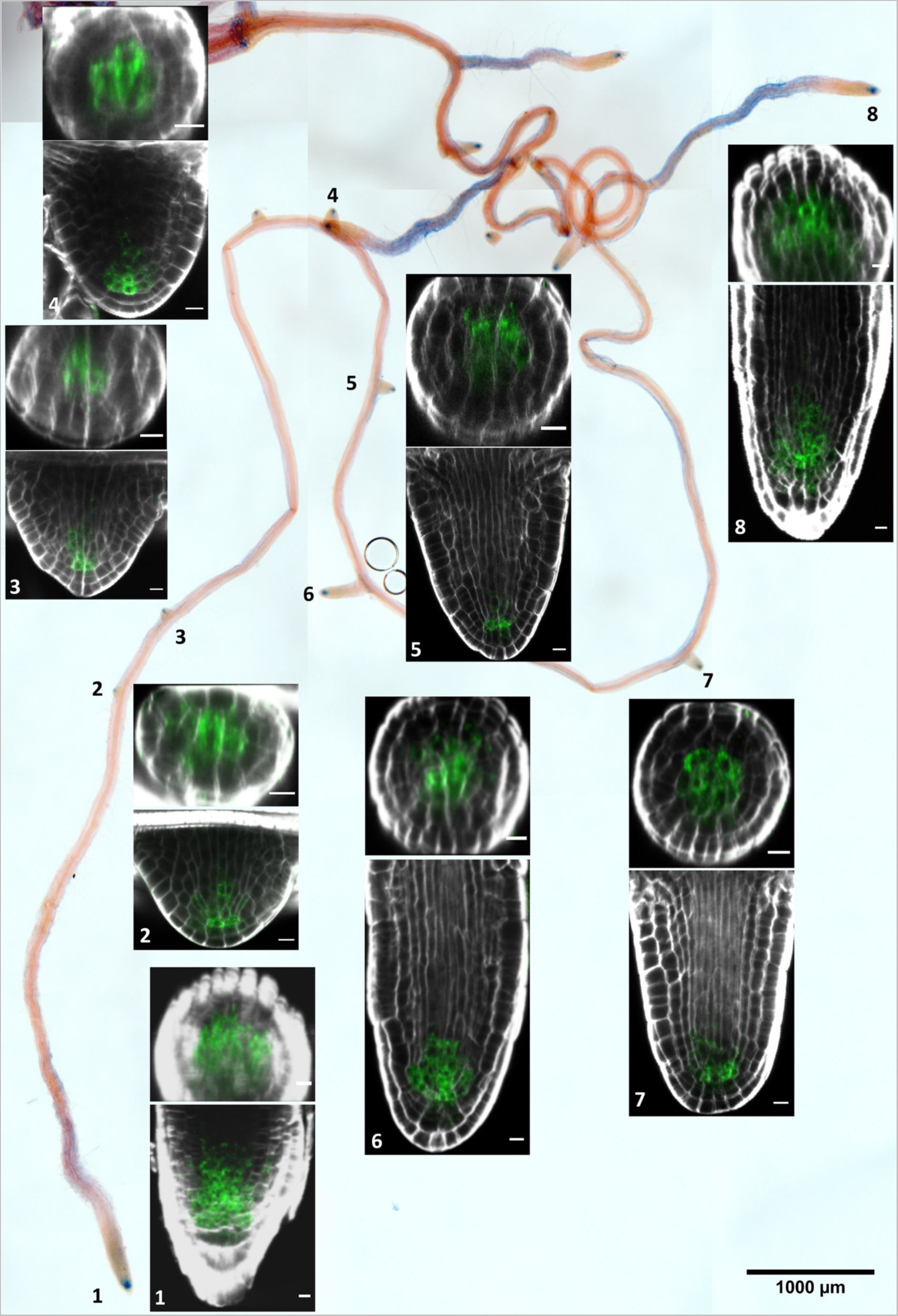
*PLT1* mRNA localization in root meristems of 10 days-old Arabidopsis seedlings. Colorimetric detection of *PLT1* mRNA is visualized by stereomicroscopy (blue). Fluorescence is visualized by confocal microscopy. For roots numbered from 1 to 8, corresponding frontal and transversal planes in confocal microscopy are presented. Cell walls stained with Direct Red 23 (red). Scale bars 1000µm for the general view, 10µm for optical sections.

### Vector® Blue Substrate Kit detection combined with tissue clearing ensure strong and uniform mRNA detection across whole organs

For probe labeling and detection, we chose a robust and cheap approach combining RNA labeling by *in vitro* transcription with DIG-UTP or FITC-UTP and detection with anti-DIG (or FITC)-antibody type Fab conjugated with Alkaline Phosphatase (AP). For AP detection we used Vector® Blue Substrate Kit that gives simultaneously colorimetric blue and fluorescent Far Red signal collected in a wavelength interval where plant tissues emit a lowest autofluorescence. The colorimetric signal allows easy monitoring the progress of the signal revelation enzymatic reaction to timely stop it. Next, the fluorescent signal can be visualized under confocal microscope with very high precision within specific tissues and individual cells and with limited autofluorescence (Figure 1, Figure S3, Figure S4). However, due to the thickness of samples, deeper detection of the signal was often reduced by fluorescence absorption and diffusion across the whole tissue. Therefore, to improve the signal, we incubated samples after detection in ClearSee solution (Kurihara *et al*., 2015). Very strong signal over the whole z stack can be observed after one week of clearing. Furthermore, samples can be stored in ClearSee for months at room temperature without signal loss.

### A standard protocol allows RNA WISH in various organs and tissues

To evaluate the versatility of our protocol, we performed hybridization on different tissues: roots, shoots, leaves and embryos of Arabidopsis grown either *in vitro* or in the greenhouse. These organs are characterized by different permeabilities and accessibilities for RNA probes that previously necessitated modifications of treatments during hybridization (Hejátko *et al*., 2006; Rozier *et al*., 2014). We selected at least two probes for each organ. The precise characterization of gene expression pattern requires counterstaining to localize signal inside tissues. For that, we combined *in situ* hybridization with a cell wall or nuclei counter staining. For the cell wall staining, we used either SR2200 dye (Musielak *et al*., 2015) that can be added to fixative (Figure 2–4), Fluorescent Brightening (FB) (Figure S1, S2, S4, S5) or Direct Red 23 that can be added to ClearSee (Figure 1, Figure S3). For nuclei staining we added DAPI to the mounting medium (Figure S6).

**Figure 2.**
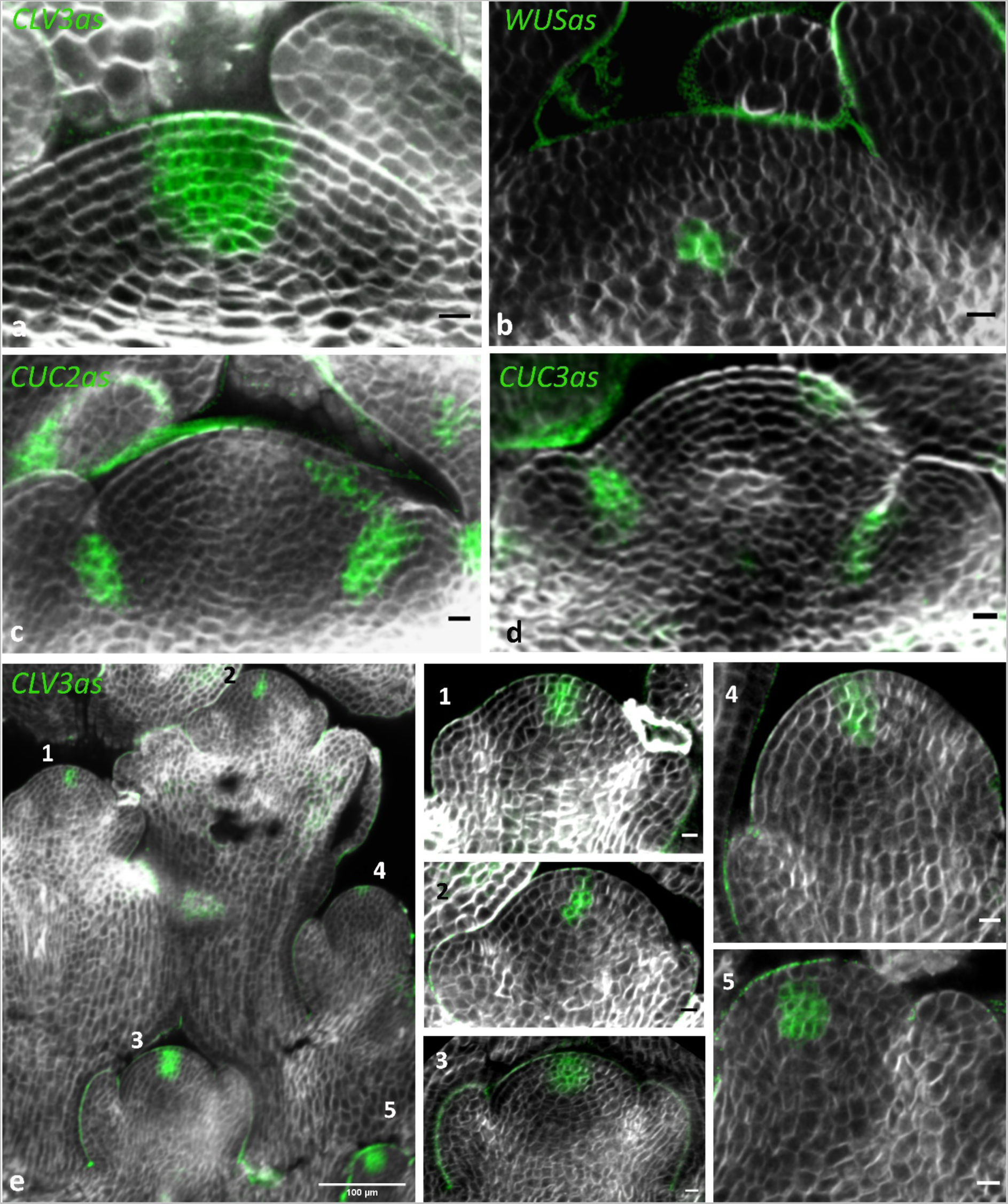
*CLV3*, *WUS* or *CUC2* mRNAs detection in SAM or axillary meristems of greenhouse-grown plants. *CLV3* (a), *WUS* (b), *CUC2* (c) expression in SAM of 4-week-old greenhouse-grown plants, *CUC2* (d). *CLV3* mRNA detection on axillary meristems of entire emerging inflorescence (e) from 6-week-old greenhouse-grown plants. Details meristems 1 to 5 (f). Asterisk indicate the absence of *CLV3* transcripts in L1. Cell walls stained with SR2200. Scale bars 10µm.

### Main root and lateral root meristems

In order to test our protocol on roots, we first analyzed in details *mRNA* transcript accumulation of *PLT1* and *WOX5* genes. To be able to count cellular layers and thereby to define developmental stages of lateral roots according to (Malamy and Benfey, 1997), we combined M2WISH with FB cell wall staining. We observed that in all protruded roots (stage VII and older) with a well-developed meristem, initials were labeled with *PLT1* and the quiescent center (QC) with *WOX5* probe respectively (Figure S5) as described in (Aida *et al*., 2004; Sarkar *et al*., 2007). *PLT1* transcripts could be detected within LR primordia from stage V onwards (Figure S5a-i) and *WOX5* transcripts could be detected as early as stage IV (Figure S5j-r). In addition, we were able to follow the evolution of *PLT1* and *WOX5* expression pattern during root development, with a progressive enlargement of *PLT1* expression domain and a progressive restriction to QC for *WOX5* transcripts. We then analyzed *HISTONE 2B* (*H2B*) and *CYCLIN B1* (*CycB1*) transcription in lateral roots combining a WISH protocol with DAPI counterstaining. Both genes are transcribed within clusters of proliferating cells (Figure S6). *CycB1* transcripts are detected concomitantly with DAPI-stained mitotic figures (Figure S6a-f), that is in agreement with a sharp peak of mitotic cyclin activity at G2/M (Colon-Carmona *et al*., 1999). In contrast to *CycB1*, *H2B* transcripts are completely excluded from cells during mitosis (DAPI), they are detected in clusters within cells bearing interphasic nuclei (Figure S6g-i) that is in accordance with their expression in meristems (Jiang *et al*., 2020).

### Shoot apical and axillary meristems

We tested our protocol by hybridizing *CLV3*, *WUS*, *CUP-SHAPED COTYLEDON 2 (CUC2)* and *3 (CUC3)* probes on shoot apical meristems of 6 weeks-old plants grown in the greenhouse. We routinely detected abundant *CLV3* transcripts in the central zone (Figure 2a), *WUS* transcripts were restricted to the organizing center (Figure 2b) and *CUC2*, *CUC3* transcripts localized to the boundary between SAM and leaf primordia (Figure 2c-d). All axillary meristems from entire apices collected shortly after bolting were marked by CLV3, demonstrating the reliability of our method (Figure 2e).

### Leaves

To extend M2WISH to other less permeable aerial organs, we performed *CUC2* and *CUC3* detection on leaves from plants growing in the greenhouse. We detected *CUC2* and *CUC3* expression in the sinus of leaf serrations as described in (Nikovics *et al*., 2006; Hasson *et al*., 2011) (Figure 3a-g, Figure S7). Interestingly, on the entire rosette, strong *CUC2* labelling could be detected on all successive leaves over a wide range of sizes (Figure S8). We tested our protocol on vascular meristematic tissue of leaves by hybridizing *PIN-FORMED 1 (PIN1*) probe. *PIN1* has a key role in leaf vascular patterning and leaf margin development as shown by the expression of a functional *pPIN1::PIN1::GFP* translational fusion in transgenics (Scarpella *et al*., 2006; Bilsborough *et al*., 2011). We hybridized entire rosettes with *PIN1* probe and obtained patterns of transcripts accumulation similar to those revealed by the translational reporter (Scarpella *et al*., 2006). *PIN1* transcripts first marked the pre-pattern of vascular network in young leaves (Figure 3h-j) and progressively disappeared in older leaves, while cells started to differentiate in a distal to proximal gradient (Figure 3k). Reconstruction of transversal sections revealed that *PIN1* is preferentially expressed at the adaxial side of the leaf vasculature (Figure 3j). *PIN1* transcripts were also detected in hydathodes, in agreement with transcriptomic analysis (Routaboul *et al*., 2022) (Figure 3o-p), and at the tip of nascent serrations at the leaf margin (Figure 3l, n) (Bilsborough *et al*., 2011). To our knowledge, this is the first time *PIN1* transcripts was detected using WISH.

**Figure 3.**
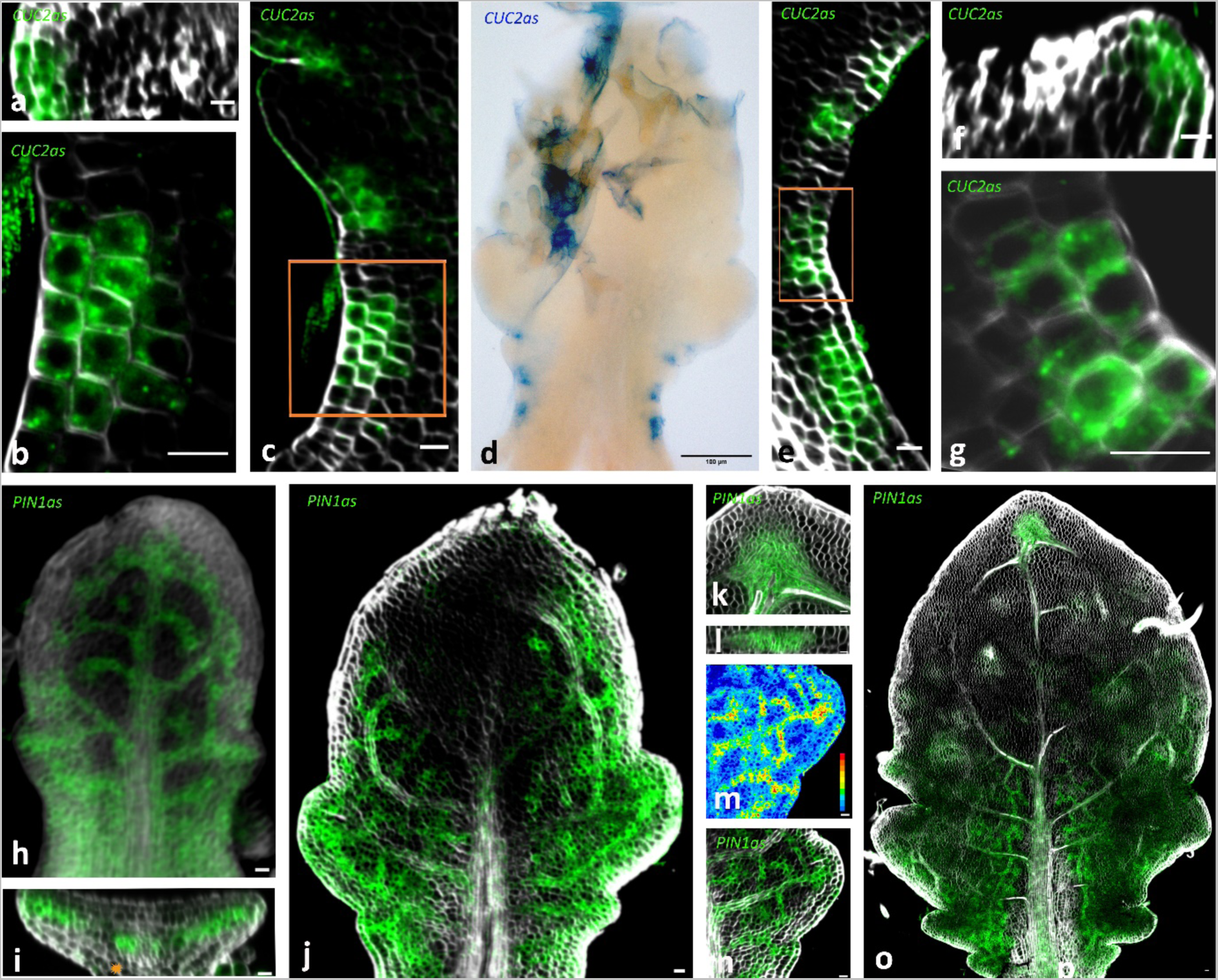
*CUC2* and *PIN1 mRNAs* detection in leaves 2-week-old greenhouse-grown plants. *CUC2 mRNA* detection in the sinus of leaf serrations (a-g). General view of *CUC2* localization (d) with a colorimetric detection. Details of left (c) and right (e) sides of the leaf (d), frontal and reconstructed transversal sections of a close-up showing details of *CUC2* transcripts accumulation (a, b-g, f). *PIN1 mRNA* detection in vascular meristematic tissue of leaves (h-o). *PIN1* transcripts mark the pre-pattern of vascular network in a young leaf (h,i). Detail of reconstructed transversal section (j) showing *PIN1* transcripts accumulation on the adaxial side of the vasculature. Asterisk indicate protophloem cells on the abaxial side. *PIN1* transcripts distribution in a distal to proximal gradient in older leaves (j,o). Detail of the proximal region of leaf (l), signal intensity shown in rainbow LUT (m). Detail of *PIN1* transcripts associated with apical hydathode (k-frontal and i-reconstructed transversal section). Cell walls stained with SR2200. Scale bars 10µm.

### Embryos

To further challenge our protocol, we tested gene expression on *Arabidopsis* embryo inside their integuments. *CUC2, SHOOT MERISTEMLESS (STM)* and *PLT1* genes were probed on seeds of different developmental stages. While heart stage and older embryos were easily dissected from seed coat after M2WISH, globular and triangular stage embryos were often damaged upon dissection. Confocal imaging of entire seed was rendered difficult due to the tannin’s precipitation in the most internal seed layer (data not shown). To overcome this problem we performed M2WISH on pale-colored seeds of *tt4-1* mutant (Burbulis *et al*., 1996). *tt4-1* is a null mutation in *TRANSPARENT TESTA* 4 coding for *CHALCONE SYNTHASE*, the first enzyme of flavonoid biosynthesis. Homozygous *tt4-1* mutants are completely devoid of flavonoids and embryos have an enhanced fatty acid biosynthesis and reduced *PIN–FORMED 4* expression, which however does not perturb embryo morphogenesis (Xuan *et al*., 2018). Using *tt4-1* mutants, we obtained patterns of *CUC2, STM* and *PLT1* transcripts accumulation from *in testa* embryos (Figure 4, Figure S9) similar to the ones previously described by (Aida *et al*., 1999; Aida *et al*., 2004; Long *et al*., 1996) on embryo sections. By analyzing the frontal, sagittal and transversal optical sections within the same sample (Figure 4, a-p and Figure S9, p-u, movie1) we could recapitulate the pattern evolution of *CUC2* and *STM* transcripts accumulation during different developmental stages. From triangular stage (transition), *CUC2* and *STM* are organized as a strip in the node of two emerging cotyledons (Figure 4a-i; Figure S9p-r). At torpedo stage, *STM* pattern becomes restricted to the central zone corresponding to future SAM (Figure S9s-u). At bent-cotyledon stage, *CUC2* transcripts progressively disappeared from the central zone and become restricted to the boundary region of cotyledon margins (Figure 4m-p). *PLT1* transcripts appeared early in the lower part of embryo (not shown) then they are detected in the QC and surrounding stem cell niche of the embryonic root (Figure S9a-o) (Aida *et al*., 2004). In summary, M2WISH protocol combined with cell wall staining gives the opportunity to determine the 3D transcripts accumulation patterns within entire embryo without laborious embryo extraction from the seed *testa*.

**Figure 4.**
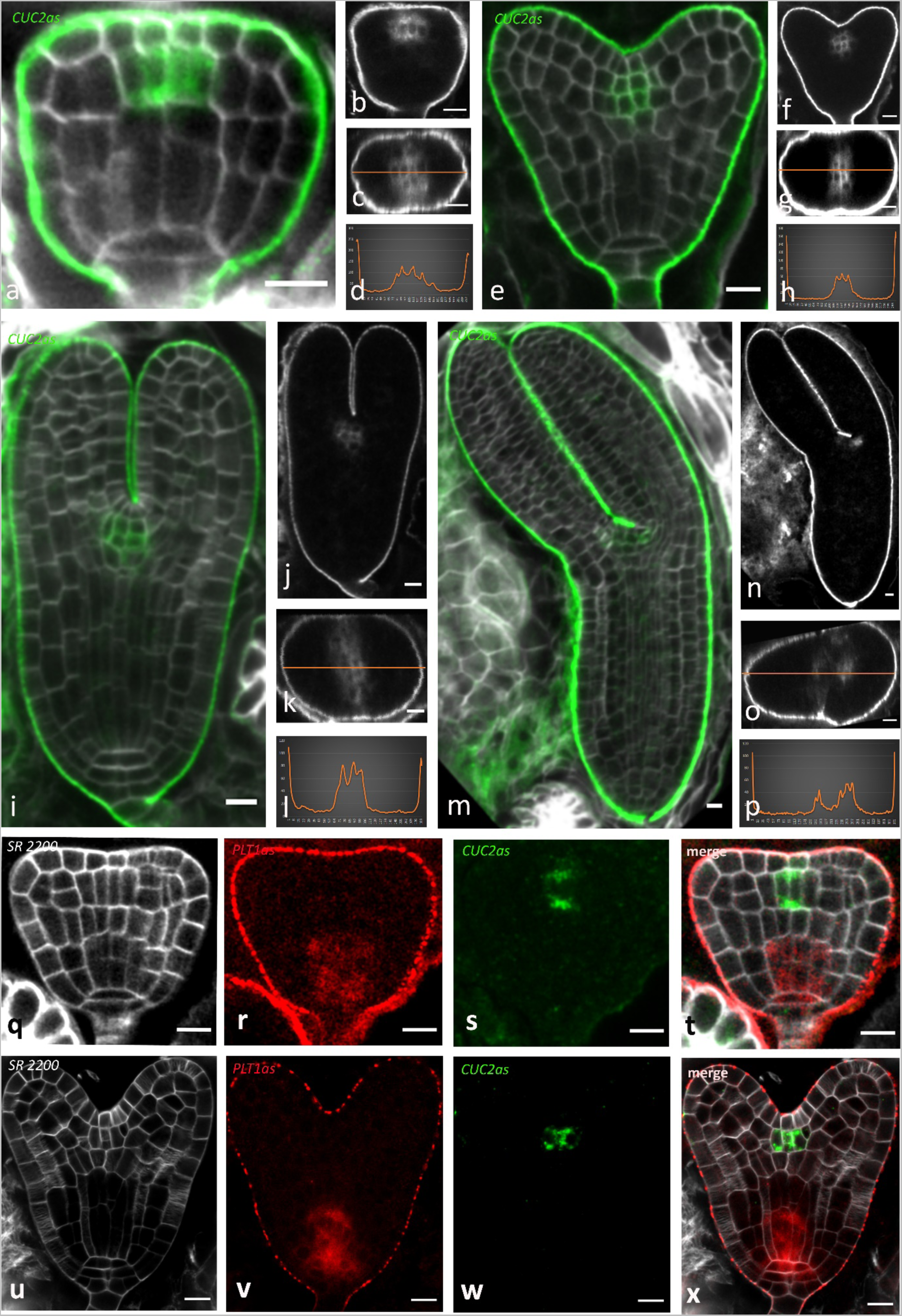
*CUC2* and *PLT1* mRNA detection in Arabidopsis embryos. Pattern evolution of *CUC2 mRNA* in *tt4* mutant (a-p). Merge images of embryo of triangular (a), heart (e), torpedo (i), walk-stick (m) stages and corresponding frontal (b, f, j, n) and reconstructed transversal optical sections (c, g, k, o). Plot of fluorescence intensity along the orange line showing homogenous signal in early stages (d, h, l) and loss of expression in the central zone of the future meristem at later stage (p). Pattern co-evolution of *PLT1* and *CUC2* transcripts accumulation in *tt4* embryo of triangular (q) and heart (b) stages. *CUC2* transcripts are detected with Vector® Blue Substrate Kit (AP activity) (a-p). *PLT1* transcripts are detected with Vector® Blue Substrate Kit (AP activity) and *CUC2* with Alexa Fluor™ 488 Tyramide Reagent (POD activity) (q-x). Cell walls stained with SR2200. Scale bars 10µm.

### Two different substrates to ensure a simultaneous detection of two genes without cross-talk

To further evaluate the versatility of our protocol we performed a co-detection of two mRNAs transcripts on meristematic tissues of embryo, shoots and roots. We produced two different riboprobes by labeling them with either digoxigenin (DIG) or fluorescein (FITC) which were simultaneously hybridized to the sample. Next, the samples were incubated with a mix of Anti-DIG antibodies conjugated to AP and Anti-FITC antibodies conjugated to POD, so different probes could be detected by different enzymatic activities. We detected consecutively POD activity applying Alexa Fluor™ 488 Tyramide Reagent and then AP activity with a Vector® Blue Substrate Kit. This double detection setup enables accurate and specific detection without the need to quench the enzymatic activity between two successive detections (Rozier *et al*., 2014), thus simplifying and providing robustness to the protocol. To confirm the co-detection specificity by double M2WISH, we simultaneously hybridized *CUC2* and *PLT1* on embryos (Figure 4, q-x). We obtained an accurate pattern of transcripts accumulation for both genes, similar to simple M2WISH (Figure 4, e, i; Figure S9, g,m). Then we tested the protocol of double M2WISH on SAM and RAM (Figure S10). We hybridized *CLV3* and *WUS* probes to the shoot apical meristem and obtained non-overlapping patterns of transcripts accumulation with *CLV3* detected in the stem cells and *WUS* in the organizing center (Figure S9, a-c). For root apical meristem, when hybridizing *PLT1* and *WOX5,* we detected *PLT1* transcripts in the initials of root meristem and in quiescent center, respectively (Figure S9, d-j). We observed equal sensitivity with both detection systems, but we noticed that AP detection gives an additional cuticula staining whatever probes or organs is used. Fortunately, the residual AP activity in cuticula has no impact on transcript accumulation detection since it occurs outside the organs.

### Expression analysis of WUS and CLV3 during meristem conversion

Meristem conversion is the process that allow a nascent Lateral Root Primordia (LRP) to be converted into SAM upon hormonal treatment (Rosspopoff et al., 2017). Meristem conversion is triggered when 5-days-old plantlets are subjected to auxin treatment for 44 hours to induce numerous divisions in the pericycle at the xylem poles (Atta *et al*., 2009) that generate lateral dividing zones, including LRP. The samples are then transferred on CK medium to induce shoot formation. We previously demonstrated that changes in gene expression profiles during LRP conversion reflects the switch in organ identity (Rosspopoff *et al*., 2017). However, cellular resolution of mRNA expression lacked in this study as well as the study of the diversity in the structures forming new shoot meristems. Therefore, we used here M2WISH to carefully analyze *WUS* and *CLV3* expression patterns during meristematic conversion induced by CK treatment in numerous samples.

After 1 day of CK treatment, *WUS* was expressed in lateral outgrowing zones corresponding to the canonical LRP composed by several cell layers depending on the LRP stages and previously described in (Rosspopoff *et al*., 2017) (Figure 5a). In addition, we detected *WUS* transcripts in flat, thickened zones composed of 3 to 5 layers of Actively Dividing Pericycle (ADP) (Figure 5c). In contrast, *WUS* expression was absent from regions where the pericycle divided less actively. *WUS* expression was induced in most LRP (41 out of 48) and in all ADP (n=6). In both LRP and ADP, we detected *WUS* transcripts within cell clusters in a central domain close to the vasculature but never extending to the L1 layer (Figure S11a). *CLV3* expression was absent in the lateral diving zone of the root following 44h auxin treatment, but present as expected on the seedling SAM (Figure S12). After 1 day on CK, *CLV3* was expressed in a minority of LRP (8 out of 65) but in all DP (n=9) (Figure 5b, d, S13). In LRP, *CLV3* was expressed in an upper domain which however remained excluded from the external layer (L1) (Figure 5b). In contrast, in ADP, *CLV3* transcripts were detected in a domain including the L1 (Figure 5d). After 2 days of CK treatment, although the proportion of *CLV3*-expressing LRP dramatically increased, *WUS* and *CLV3* expression patterns remained unchanged (Figure S11b; Figure S13, b). At 3 days of CK treatment, the LRP dome flattened as described previously by (Rosspopoff *et al*., 2017), prefiguring an early promeristem (Figure 5e,f,h). At this stage, it was difficult to morphologically distinguished between flattened structures coming from LRP and those coming from developing ADP. Nevertheless, we still observed two different *CLV3* gene expression patterns: 1) a sub-epidermal *CLV3* pattern (Figure 5f) and 2) a *CLV3* pattern that included the L1 (Figure 5h). Finally, at 5 days of CK treatment, we observed canonical shoot meristems with a dome and two or more leaf primordia (Figure 5i,j; Figure S11c and Figure S13c). At this stage a unique pattern of *CLV3*/*WUS* was observed, with *CLV3* being expressed in an apical domain extending into the L1, whereas *WUS* expression was mostly confined to a sub-epidermal zone corresponding likely to the organizing center. Taken together, these results suggest that SAMs result from the convergence of two different structures that differ in their initial shapes and *CLV3* expression patterns.

**Figure 5.**
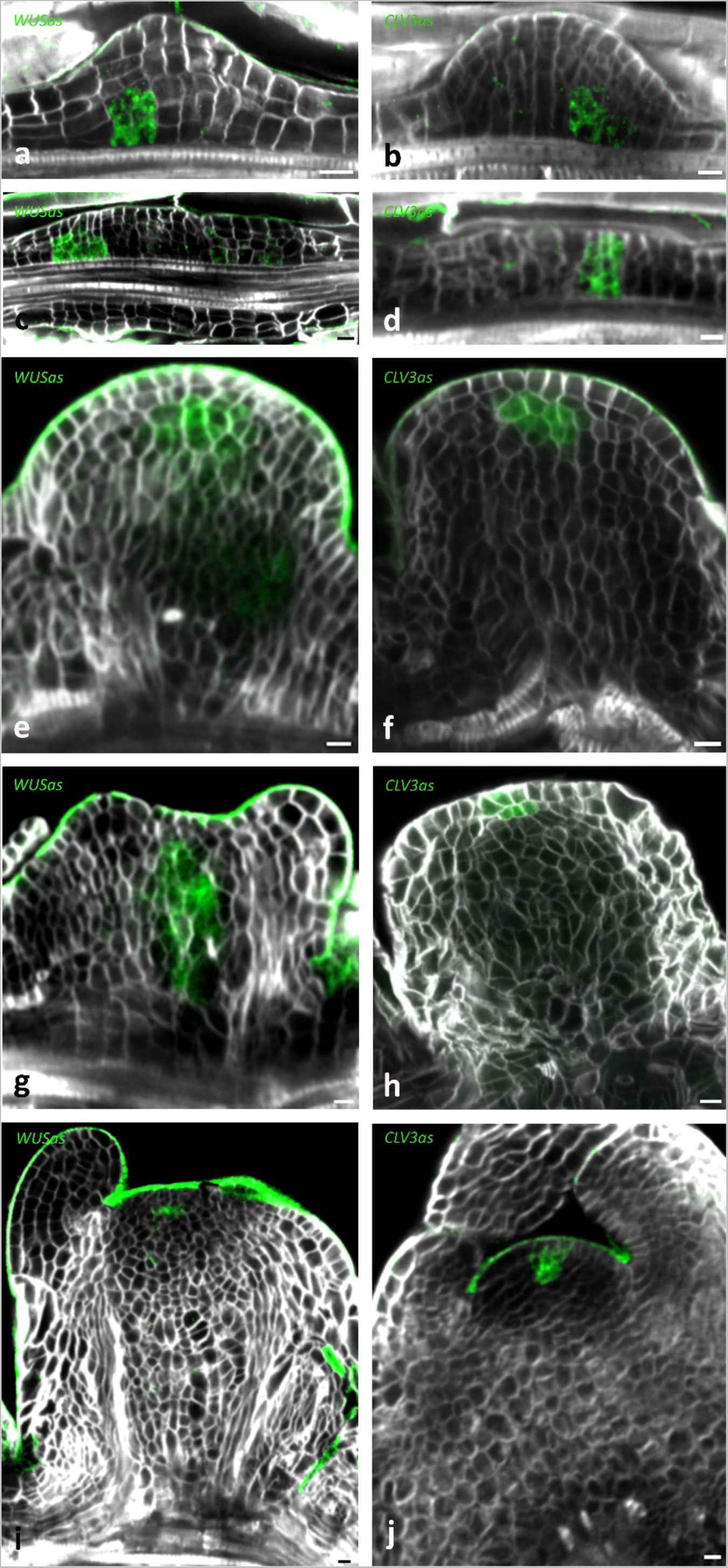
Representative *WUS* and *CLV3* gene expression patterns during different stages of meristem conversion. LRP stage(a-b), ADP stage (c-d), early promeristem (e-h), SAM (i-j). *mRNAs* transcripts are detected with Vector® Blue Substrate Kit (AP activity). Cell walls stained with SR2200. Scale bars 10µm.

Because the comparative analysis of *WUS* and *CLV3* patterns established by single WISH is limited by the level of the synchrony between individual lateral zones, we applied a double WISH and analyzed *WUS* and *CLV3* co-expression patterns within the same zones on entire explants. As expected, we confirmed the previous evolution of shape and gene expression patterns from 2 to 3 days post-CK treatment observed in simple localization (Figure 6). At 2 days, 77% of *WUS* positive zones express *CLV3* (n=104), at 3 days, 89 % of converting zones express *WUS* and *CLV3* (n=65). All organs expressing *CLV3* also expressed *WUS*, in agreement with the role of *WUS* as an inducer of *CLV3* in root to shoot conversion system (Fuchs and Lohmann, 2020).

**Figure 6.**
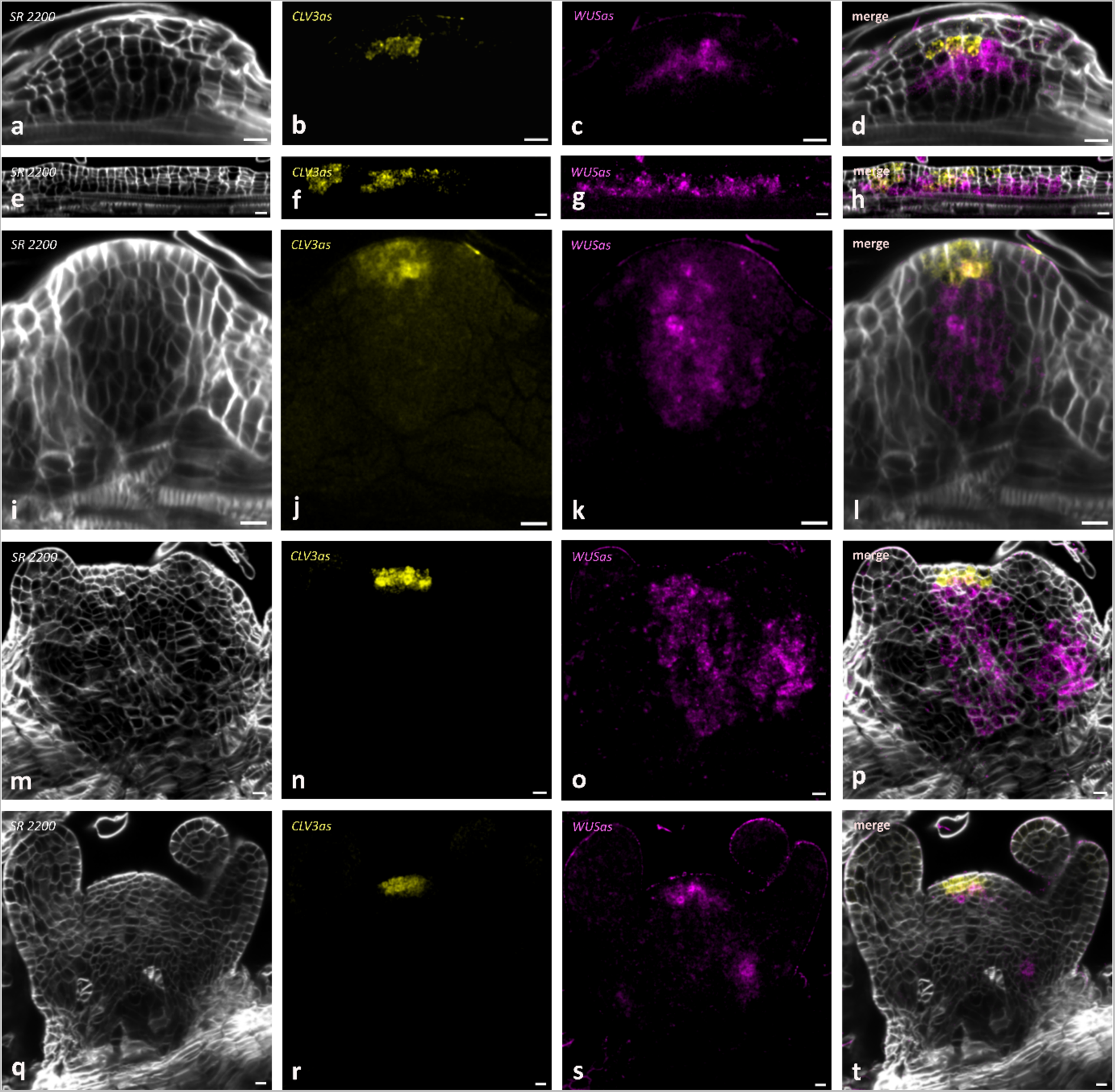
Co-detection of *WUS* and *CLV3* mRNA during different stages of meristem conversion. LRP stage (a-d), ADP stage (e-h), early promeristem (i-l), late promeristem (m-p), SAM (q-t). *WUS* mRNAs transcripts are detected with Vector® Blue Substrate Kit (AP activity in magenta) and *CLV3* mRNAs with Alexa Fluor™ 488 Tyramide Reagent (POD activity in yellow). Cell walls stained with SR2200 (gray). Scale bars 10µm.

### 3D map of CLV3 and WUS gene expression in converting organ at a cell resolution

In order to assess the variability of *CLV3* and *WUS* expression within organs undergoing conversion, we created a 3D representation of corresponding expression domains. For this we developed a pipeline taking as an input a 3D confocal image with 3 channels, corresponding respectively to cell wall, *CLV3* and *WUS* fluorescent signal (Fig. S14A). The output is a 3D map at cellular resolution representing the regions of respective gene expression domains (Fig. S14E). First, we preprocessed input images to emphasize significant signals and reduce the noise (Biot *et al*., 2008) (Fig. S14B). Next, the cell wall channel was used to segment (i.e., to identify) individual cells (Fig. S14C). For this, PlantSeg software (Wolny *et al*., 2020) was used to predict cell wall positions, then a h-watershed algorithm was applied to label individual cells (Biot *et al*., 2016). From this image, we quantified cell volume of all cells and discarded labels resulting from over-segmented small volumes. Remaining cell labels were then associated with the corresponding signal in *CLV3* and *WUS* channels (Fig. S14D). For this, we quantified the average signal values corresponding to each label. Using ground-truth expertise, a threshold was manually defined to determine regions with significant signal levels, which correspond to true expression domains. Finally, 3D surface representations were then generated to visualize these domains directly within 3D organ volumes (Fig. S14E).

This workflow was applied on 18 LRP/ADP/SAM structures from 2 to 5 days after CK treatment. In a 2-days old representative ADP structure, the 146 cells expressing *WUS* formed an elongated domain in a sub-epidermal position. The 147 cells expressing *CLV3*-formed a large zone above and adjacent to the *WUS*-expressing cells. 67 cells expressed both *CLV3* and *WUS* and we unambiguously located 29 cells expressing *CLV3* in L1 versus 3 L1 *WUS*-expressing cells (Figure 7a-i). In a 2 days-old representative LRP structure 58 *WUS*-expressing cells were present in a central position in the dome, whereas 22 *CLV3*-expressing cells occupied a sub-epidermal zone above and adjacent to the *WUS* domain. Only 2 cells expressed both *WUS* and *CLV3* and no cell expressing *WUS* or *CLV3* was found in the L1 (epidermis layer) (Figure 7j-q). At 5 days for CK treatment, the *CLV3* and *WUS*-expressing cells were present in zones corresponding to a canonical shoot apical meristem i.e. a meristem dome surrounded by several leaf primordia (Figure 7r-y). The *WUS* expression domain was composed of 310 cells spreading over the entire meristem except the L1. The 107 cells expressing *CLV3* were restricted to a central zone and included 31 cells located in the L1. We found 26 cells expressing both *CLV3* and *WUS*.

**Figure 7.**
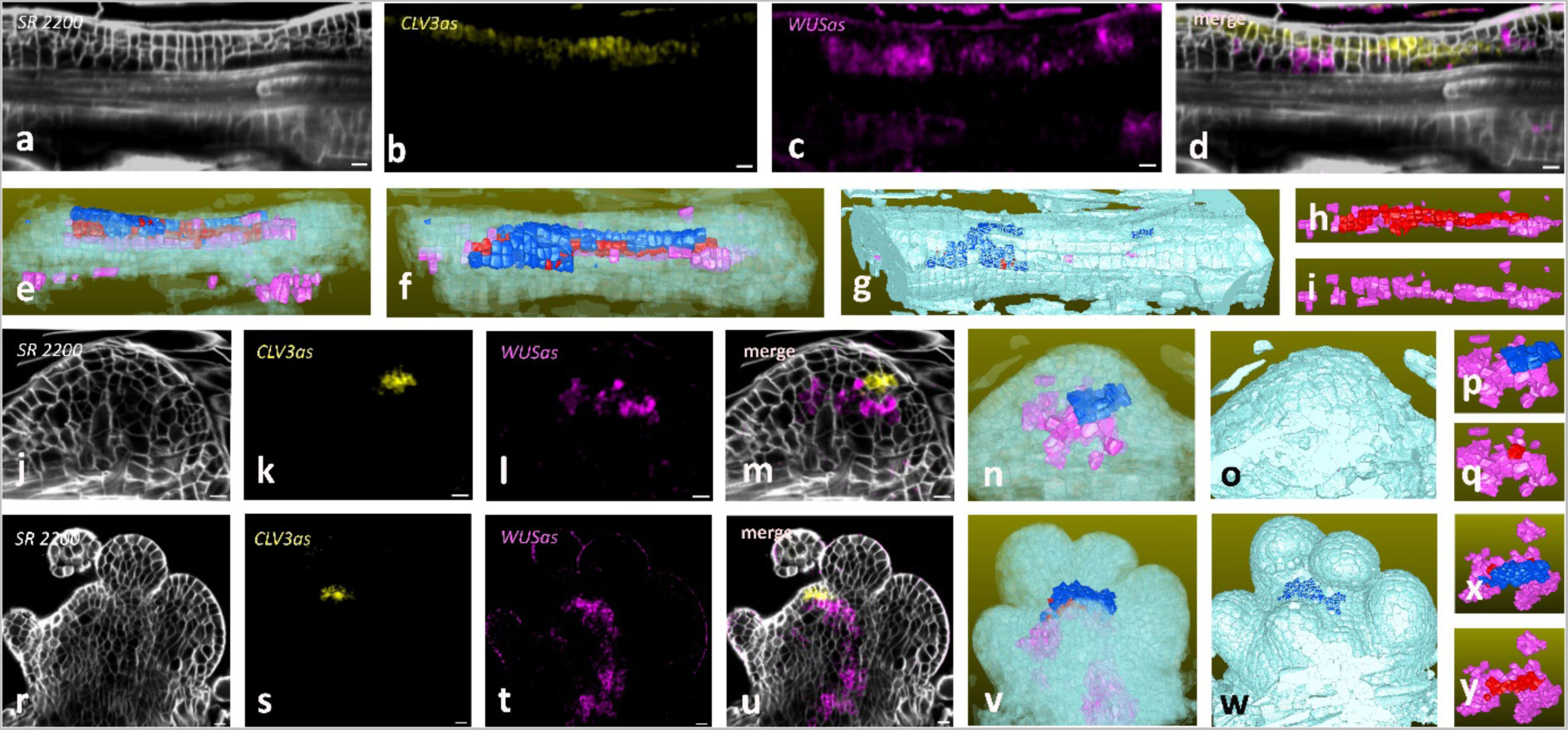
3D representation of *WUS* and *CLV3* gene expression patterns during meristem conversion. mRNA detection of *CLV3* and *WUS* mRNAs in ADP structure (a-d), LRP structure (j-m), SAM (r-u). 3D representation of the corresponding ADP structure (e-i), LRP structure (n-q), SAM (v-y). Cells are color-coded: pink (*WUS*-expressing cells), blue (*CLV3*-expressing cells), red (*WUS*/*CLV3* co-expressing cells), pale blue (other cells). Cell walls stained with SR2200. Scale bars 10µm.

Using this 3D representation, we were able to unambiguously identify cells expressing *WUS* and/or *CLV3* within the wider organ context. We confirmed our previous observations that meristem conversion can occur from different structures, LRP and ADP, which later development converges into a SAM. We also show that early *CLV3* patterns differ between these two structures being either present in the L1 layer in ADP structures or confined to sub-epidermal layers in LRP structures. These differences progressively faded over the time-course of conversion to give rise to a more canonical SAM pattern. Together, our analysis reveals a high flexibility and variability in the process of meristem conversion with different developmental paths leading to identical final organization of established meristems.

## Discussion/Conclusion

Here, we proposed a new M2WISH protocol that can accommodate for Arabidopsis organs with different permeability. The combination of MW treatment, clearing, and counter-staining ensure an efficient detection of gene expression with an appropriate cell resolution. This is also a method of choice for the characterization of expression patterns in complex genetic backgrounds, such as natural accessions or recombinant inbred lines that would necessitate time-consuming crossing and selection to use transgenic reporter constructs. The detailed protocol with precise step-by-step instructions can be found in Additional file 2.

Using a cell resolution, we showed a 3D representation of *CLV3/WUS* gene expression pattern co-evolution. We had previously used this protocol to study *CLV3/WUS* gene expression pattern dynamics in *de novo* stem cell formation during the formation of Axillary Meristem (AM) (Nicolas *et al*., 2022). Furthermore, *CLV3/WUS* gene expression pattern dynamics is subjected variations depending on whether rosette or cauline AM are formed (Nicolas and Laufs, 2022). In rosette AM, an initially overlapping central region of *WUS/CLV3* expression is progressively resolved into a *CLV3* domain in L1 and L2 and a lower *WUS* domain (Xin *et al*., 2017). On the other hand, in cauline AM both *WUS* and *CLV3* are present in a initially apical domain while later *WUS* operates a dynamic shift to the central domain (Nicolas *et al*., 2022). In this study, we provided evidences that *WUS* always precedes *CLV3* expression. Then, *CLV3* expression localized either on the L1 and subepidermal layers for ADP structures, or in a central domain of LRP structures and then moves into an apical domain at later stages. Such a variation suggests that structures prone to convert have different origin and initial organization. Indeed, competent auxin-mediated LRP were shown to express both WOX5 and PLT1 in a central domain (Rosspopoff et al., 2017) as in indirect regeneration through Callus Induced Medium point where WOX5-expressing middle cell layer acts as a key player for pluripotency acquisition and organ regeneration (Zhai and Xu, 2021). Whether *WOX5* and *PLT1* are expressed during ADP structure conversion remains to be determined.

Altogether, M2WISH provides a reliable and robust method to systematically validate the expression of new genes with high precision and thus is an adapted tool to validate results obtained with new technics of single cell transcriptomics

## Supporting information

detailed protocol

## Acknowledgments

The IJPB benefits from the support of Saclay Plant Sciences-SPS (ANR-17-EUR-0007). We are grateful to A. Berger for her support. This work has benefited from the support of IJPB’s Plant Observatory technological platforms. The authors thank J.Burguet and N. Arnaud for critical reading of the manuscript

## Author contributions

LC, PL and JCP conceptualization; LC, HM, AN, PR, MdC, LB and JCP performing the experiment; LC, EB and JCP post-processing imaging; LC, EB, PL and JCP writing the manuscript; all authors edited and reviewed the mnuscript

## Materials & Methods

### Probe synthesis and labelling with in vitro transcription

RNA labelled probes were prepared with *in vitro* transcription of PCR amplified fragments from cloned cDNA. For antisense probes, a T7 promoter sequence (TGTAATACGACTCACTATAGGG) was added to 3’ gene-specific primers (Additional file1. The detailed protocol). For labelling of 1µg DNA template, the mix containing T7 RNA Polymerase (Promega, Cat No: P2075), Digoxigenin-11-UTP (Roche, Cat No: 11209256910) or Fluorescein-12-UTP (Roche, Cat No: 11373242910) was prepared (Additional file1). RNA probes were then hydrolyzed into fragments of 100 bp on average before hybridization. To estimate the labelling efficiency, dilution series of DIG/F-labelled probes were blotted on a positively charged nylon membrane and detected for Dig with Anti-Digoxigenin-Alkaline phosphatase (AP), Fab fragment (Roche 11093274910) and for Fluorescein with Anti-Fluorescein-AP, Fab fragments (Roche, Cat No: 11426338910). AP activity was visualized with Vector® Blue Alkaline Phosphatase Substrate (Vector Laboratories, SK-5300) according to supplied protocol. For two probes detection, DIG-labelled probes were detected with Anti-Digoxigenin-POD, Fab fragments (Roche, Cat No: 11207733910) and visualized with Vector® VIP Substrate Kit, Peroxidase (HRP) (Vector Laboratories, Cat No: SK-4600) according to supplied protocol.

### Sample fixation and permeabilization

Samples were fixed in 4% paraformaldehyde (PFA) (Sigma-Aldrich, P6148), 0.1% Triton X-100 (Sigma-Aldrich, Cat No: T9284), 0.1% SR2200 Cell Wall Stain (Renaissance Chemicals) for 1 hour under vacuum and stored at 4°C for 24 hours. Next, they were dehydrated in a gradient of ethanol concentrations going from 10 to 70%. For each concentration, samples were microwaved 5 times for 30 seconds at 180 W. After this step, samples can be stored for up to 2 months at −20°C in ethanol 70%.

Samples were then rehydrated and treated with Proteinase K 10µg.ml^−1^ (Roche, Cat No: 3115836001) for 10 min at room temperature (RT), Glycine 0.2% (Sigma-Aldrich, Cat No: G7126) for 2min at RT, and finally with a mix 1.5 % Triethanolamine (Sigma-Aldrich, Cat No: 90279), 5% Acetic anhydride (Sigma-Aldrich, Cat No: 320102) and 0.1% HCl (Sigma-Aldrich, Cat No: H1758) for 10 min at RT. Post-fixation was done in RCL2 for 15min (EXILONE, R 1505J).

### Hybridization

Pre-hybridization was done in 50% Formamide (EUROBIO, GHYFOR01-01), 5xSSC (EUROBIO, GHYSSC00-07), 0.1% Tween-20 (Sigma-Aldrich, Cat No: 8.22184). Probes were denatured at 80°C for 2min and put on ice. For the hybridization, probes were added to 24 wells tissue culture plate (VWR, 734-2325) containing samples in a hybridizing buffer (50% Formamide (Eurobio Scientific, cat No: GHYFORO1-01), 10% Dextran Sulphate (VWR Chemicals, Cat No: 0198), 0.5mg.ml^−1^ tRNA (ROCHE 10109509001), 0.01% Tween-20, 2.5X Denhardt and left overnight at 47°C.

### Washouts

After hybridization samples were incubated in 0.1xSSC (Eurobio Scientific, Cat No: GHYSSC00-07), 0.5% SDS (Eurobio Scientific, Cat No: GHYSDS02) for 30min at 47°C, then in 2xSSC, 50% Formamide for 60min at 47°C. In order to remove non-hybridized single strand RNA, samples were buffered for 5min in TNE (10mM Tris (Eurobio Scientific, Cat No: GAUTRI00-66), 1mM EDTA (Sigma-Aldrich, Cat No: E3889), 100mM NaCl (Sigma-Aldrich, Cat No: S3014), pH 8.0) at 47°C, then treated with 20µg.ml^−1^ RNase A (Sigma, Cat No: R6513) (in TNE) at 37°C during 30min and washed again for 5min in TNE at 47°C. Then samples were incubated in 2xSSC, 50% Formamide for 60min at 47°C, in 0.1xSSC for 2min at 47°C and in 1x PBS (Eurobio Scientific, Cat No: GAUPBS00-01) for 15min at RT or over night (O/N).

### Sample digestion

Samples were digested using a cocktail of enzymes (0.08% macerozyme R10 (Duchefa, M8002.0005), 0.08% cellulase RS (Duchefa, C8003.0005), 0.04% Pectolyase Y23 (Duchefa, P9004.0001), 0.12% pectinase (Serva, 31660) for 10min at RT. Then samples were rinsed 3 times in 1xPBS at RT.

### Signal detection

Signal detection was performed at RT with a gentle shaking.

For one probe detection, samples were incubated in 10mM Tris-HCl pH7.5; 15mM NaCl, 1% BSA, 0.5% Triton X-100 for 60min. Immunodetection of DIG signal was performed with anti-DIG-AP Fab antibodies 1:1250 dilution in 10mM Tris-Cl pH7.5; 15mM NaCl, 1% BSA, 0.5% Triton X-100 for O/N. Then samples were washed in 100mM Tris-HCl pH8,2; 0.1% Tween-2020 three times for 10min at RT. Dual colorimetric and fluorescence detection of AP activity was made with Vector® Blue Alkaline Phosphatase Substrate (Vector Laboratories, SK-5300) according to supplied protocol.

Samples then were incubated in ClearSee (10% Xylitol (Sigma-Aldrich, Cat No: W507630); 15% Sodium deoxycholate (Sigma-Aldrich, Cat No: D6750); 25% Urea (Sigma-Aldrich, Cat No: 9510-OP)) for at least 5 days. Then samples were mounted in ClearSee and visualized under confocal microscope.

For two probes detection, samples were incubated with Anti-Dig mouse monoclonal antibody (Sigma-Aldrich, Cat No: D8156) 1:250 dilution in 10mM Tris-HCl pH7.5; 15mM NaCl, 1% BSA, 0.5% Triton X-100 for O/N at 4°C then washed three times in PBS-Triton 1% for 15min at RT. Samples were Incubated with Goat-anti-mouse-POD (1/1000 dilution) and Anti-Fluorescein-Fab-AP (1/1000 dilution) in 10mM Tris-HCl pH7.5; 15mM NaCl, 1% BSA, 0.5% Triton X-100 for O/N at 4°C and washed three times in PBS-Triton 0.1% for 15min at RT. The fluorescent POD staining reaction was performed with Tyramide Signal Amplification Kit (Fisher Scientific, Cat No: 15611892) according to (Rozier et al., 2014). The AP staining reaction was performed with Vector® Blue Substrate Kit according to manufactured protocol.

Samples were washed three times in 100mM Tris-HCl pH8.2; 0.1% Tween-2020 for 10min at RT and kept in 1xPBS. One of cell wall dyes can be added to samples in PBS (Fluorescent Brightening to 50µg.ml^−1^ dilution, Direct Red 80 to 0.01% dilution, SR2200 to 1/1000 dilution). Incubate O/N at RT with a gentle shaking. Samples were mounted in Citifluor AF1(Agar Scientific, Cat No: AGR1320)/PBS solution (3:1) with (optional a DAPI (Roche, Cat No: 10236276001) to a final concentration of 10µg.ml^−1^) and proceed with confocal microscope analysis.

### Confocal microscopy and image analysis

Images were taken with a Leica SP5 confocal laser scanning microscope (Leica Microsystems). Alexa Fluor 488 excitation was excited at 488nm and its emission was collected at 500 to 550nm; Vector® Blue Substrate excitation was done at 633 nm; its emission was collected at 740 to 800nm. SR2200 was excited with a 405nm UV laser and its emission recorded between 420 and 480nm; similar settings were used to detect DAPI. Direct Red 23 was excited with 561nm laser and detected at 580 to 615nm.

For all images, 3D Gaussian blur filter was applied and orthogonal sections were produced using Fiji software. Figures were arranged in AdobePhotoshop.Cs. 3D rendering and movie export were made with OsiriX MD software.

### 3D parametric representation

The voxels in the raw image are resized by interpolation to obtain images with isotropic spatial calibration. A Gaussian filter of radius 2 followed by an attenuation correction (Biot *et al*., 2008) is then applied to all the images. Cell boundaries are processed by a convolutional neural network called PlantSeg (Wolny *et al*., 2020) to enhance the boundary signals in the input images, then segmented by a watershed to label each cell. From this segmentation, morphometric data (cell volume) is curated to eliminate under- or over-segmented cells. The *CLV3* and *WUS* images are used to create a parametric map of average fluorescence signal intensities from each cell wall channel image. This parametric map is thresholded to obtain only cells with a strong signal. Labelled cells were extracted using triangular grids corresponding to the cell boundaries. These maps were visualized using Sviewer software (https://free-d.versailles.inrae.fr/html/freed.html). All analyses were performed using BIP software (http://free-d.versailles.inra.fr/html/bip.html).

**Figure S1. *PLT1* mRNA localization in root meristems of non-MW-treated 10 days-old Arabidopsis seedlings.**

Colorimetric detection is visualized by stereomicroscopy. Fluorescence detection is visualized by confocal microscopy. For roots 1-6, frontal and transversal planes are presented. Only a part of non-MW-treated roots (1, 3, 6) bore a signal. Cell walls are counterstained with Fluorescent Brightening (FB). Scale bars 1000µm for the general view, 10µm for optical sections.

**Figure S2. *PLT1* sense probe hybridization in root meristems of 10 days-old *Arabidopsis* seedlings.**

No labelling inside root meristems is present, but unspecific staining (ex: root 1). Colorimetric detection is visualized by stereomicroscopy. Fluorescence detection is visualized by confocal microscopy. Cell walls stained with Fluorescent Brightening. Scale bars 1000µm for the general view, 10µm for optical sections.

**Figure S3. *WOX5 mRNA* localization in organizing centers of 10 days-old Arabidopsis seedlings.**

Colorimetric detection is visualized by stereomicroscopy. Fluorescence detection is visualized by confocal microscopy. For roots 1-4, frontal and transversal planes are presented. Cell walls stained with Direct Red 23. Scale bars 1000µm for the general view, 10µm for optical sections.

**Figure S4. *WUS* mRNA detection in SAM of 10 days-old Arabidopsis seedlings.**

Colorimetric detection is visualized by stereomicroscopy (a). Fluorescence detection is visualized by confocal microscopy (b). Detail of the organizing center showing accurate mRNA detection within individual cells. Asterisks indicate individual cells on this optical section. Cell walls stained with Fluorescent Brightening. Scale bars 10µm.

**Figure S5. *PLT1 and WOX5 mRN*A detection in lateral root meristems.**

*PLT1 mRNA* detection in lateral root stage V (a-c), stage VI (d-f) and stage VIII (g-i). *WOX5 mRNA* detection in lateral root stage IV (j-l) and in QC of main root (m, n, o - reconstructed transversal optical sections) and (p, q, r - reconstructed frontal optical section). Cells walls stained with SR2200. Scale bars 10µm.

**Figure S6. *CycB1* and *H2B mRNA* detection in lateral roots**

*CycB1* mRNA detection (a-f), *H2B* mRNA detection(g-i). Red asterisks indicate DAPI stained mitotic figures. White asterisks indicate concomitant detection of *CycB1* transcripts with dividing cells. *H2B* transcripts are completely excluded from cells in mitosis, they are detected in clusters of cells bearing interphasic nuclei (arrow). Nuclei stained with DAPI. Scale bars 10µm.

**Figure S7. *CUC2* mRNA detection in the sinus of leaves.**

General view of *CUC2* localization (d) with a colorimetric detection. Details of left (a-c) and right (e-g) sides of the leaf. Cell walls stained with SR2200. Scale bars 10µm.

**Figure S8. *CUC2* transcripts localization on leaves from one rosette.**

For every leaf, details of its left and right sides are shown. Closer to general view mRNAs signal, then merged with cell wall staining. Cell walls stained with SR2200. Scale bars 10µm.

**Figure S9. *PLT1* and *STM* mRNAs detection in *tt4* embryos**

*PLT1* detection in *tt4* embryos (a-o). Frontal, reconstructed transversal and sagittal sections of globular (a-c), triangular (d-f), heart (g-i), torpedo (g-o) stages. *STM* detection in *tt4* embryos (p-o). Frontal, reconstructed transversal and sagittal section of heart (p-r) and torpedo (s-u) stages. Cell walls stained with SR2200. Scale bars 10µm.

**Figure S10. Co-detection of *CLV3* and *WUS* mRNA in SAM and of *WOX5* and *PLT1* mRNAs in RAM**

(a-c), 3D reconstruction of merge image(c). *CLV3* detected with Alexa Fluor™ 488 Tyramide Reagent (POD activity), *WUS* transcripts detected with Vector® Blue Substrate Kit (AP activity). Co-detection of *PLT1* and *WOX5* transcripts on RAM (d-j). *WOX5* transcripts detected with Vector® Blue Substrate Kit (AP activity) and *PLT1* detected with Alexa Fluor™ 488 Tyramide Reagent (POD activity). Cell walls are stained with FB. Scale bars 10µm.

**Figure S11: Statistical evolution of *WUS* mRNA detection in converting roots.**

mRNA detection was performed after 1 day (a), 2 days (b) or 3 days (c) of CK treatment. Colorimetric detection is visualized by stereomicroscopy (blue). Exhaustive fluorescence detection of *WUS* mRNA is visualized by confocal microscopy. For all numbered outgrowing zone, corresponding frontal planes in confocal microscopy are presented. Pie-charts show the percentage of lateral outgrowing zones expressing *WUS* at 1, 2 or 3 days of CK treatment. Cell walls are counterstained with SR2200. Scale bars 1000µm for the general view, 10µm for optical sections.

**Figure S12.*CLV3* mRNA localization in SAM and root meristems of 5 days-old seedling treated 44h with auxin.**

Colorimetric detection of *CLV3* mRNA is visualized by stereomicroscopy (blue). Fluorescence detection is visualized by confocal microscopy. Details (a-d) of shoot meristem (e). For roots 1-4 corresponding frontal planes in confocal microscopy are presented. Cell walls stained with SR2200. Scale bar 1000µm for the general view, 10µm for optical sections.

**Figure S13. Statistical evolution of *CLV3* mRNA detection in converting roots.**

mRNA detection was performed after 1 day (a), 2 days (b) or 3 days (c) of CK treatment. Colorimetric detection is visualized by stereomicroscopy (blue). Exhaustive fluorescence detection of *CLV3* mRNA is visualized by confocal microscopy. For all numbered outgrowing zone, corresponding frontal planes in confocal microscopy are presented. Pie-charts showing the percentage of lateral outgrowing zones expressing *CLV3* at 1, 2 or 3 days of CK treatment. Cell walls are counterstained with SR2200. Scale bars 1000µm for the general view, 10µm for optical sections.

**Figure S14. Workflow analysis for a 3D representation of *CLV3* and *WUS* gene expression pattern in a converting organ.**

Multichannels confocal acquisition (A) Preprocessing (B) Cells labelling (C) Fluorescence assignment (D), 3D Visualization (E)

