## Supplemental figures for "M2WISH: an easy and efficient protocol for whole-mount mRNA *in situ* hybridization that allows 3D cell resolution of gene expression in *Arabidopsis thaliana*"

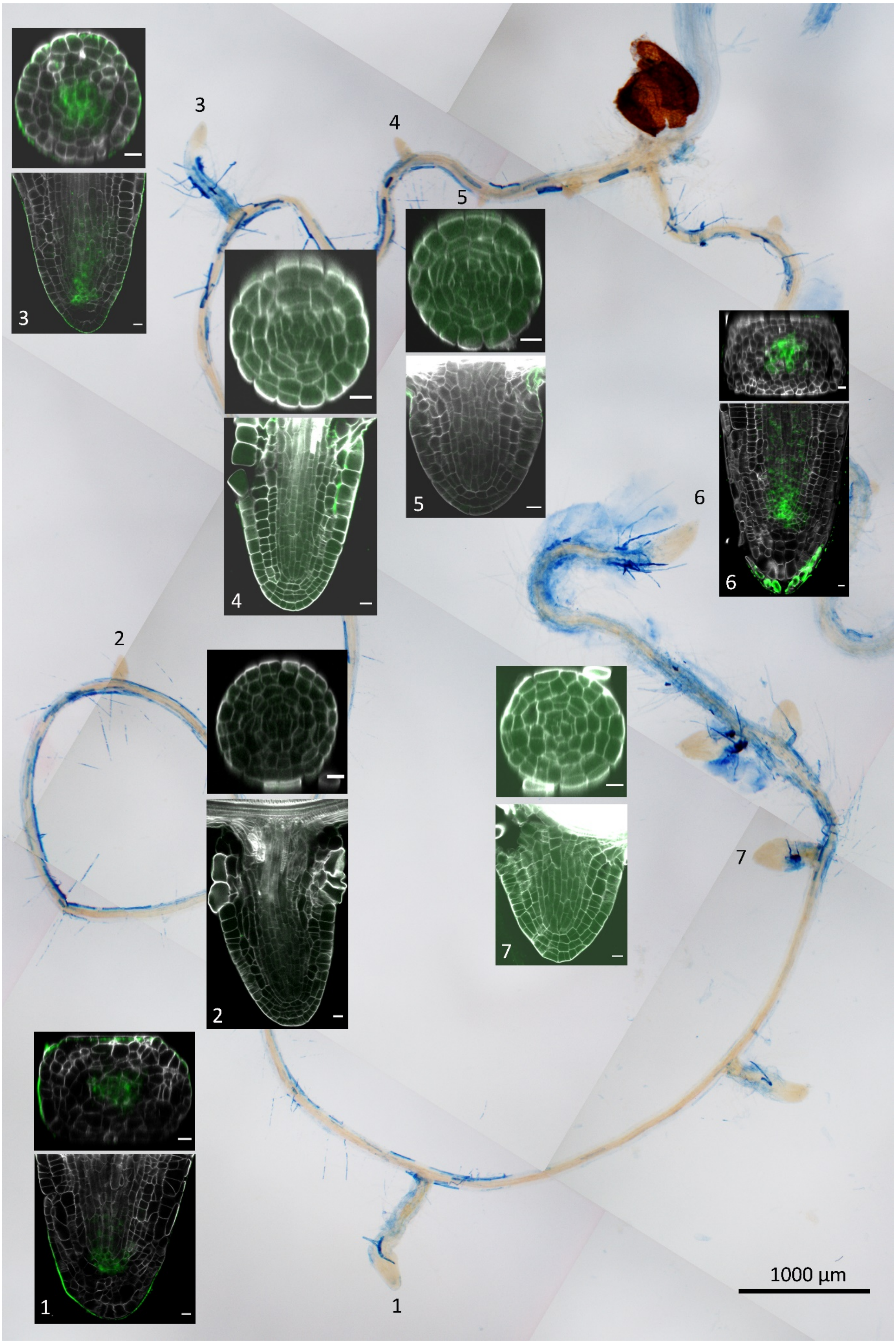


Figure S1

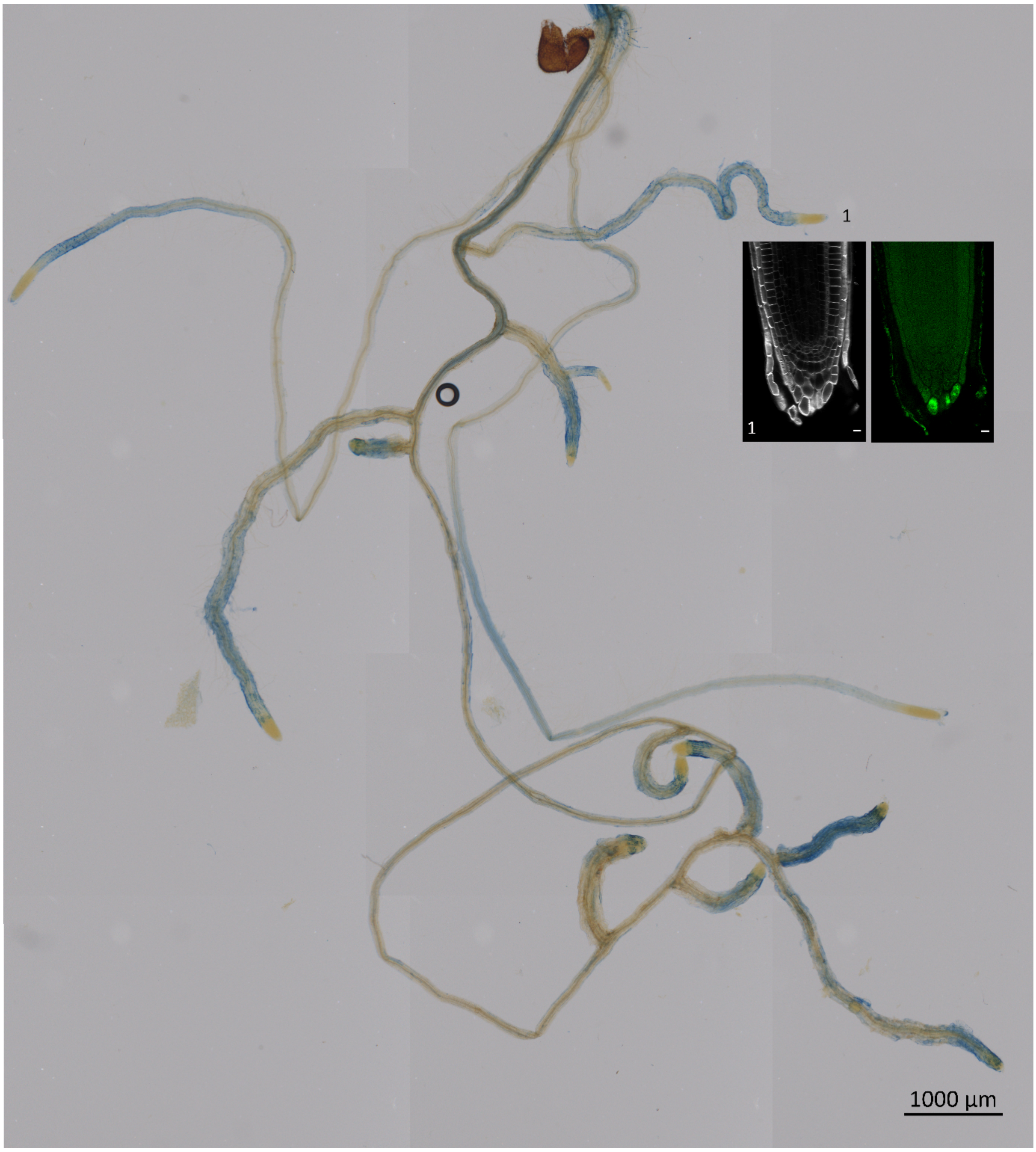


Figure S2

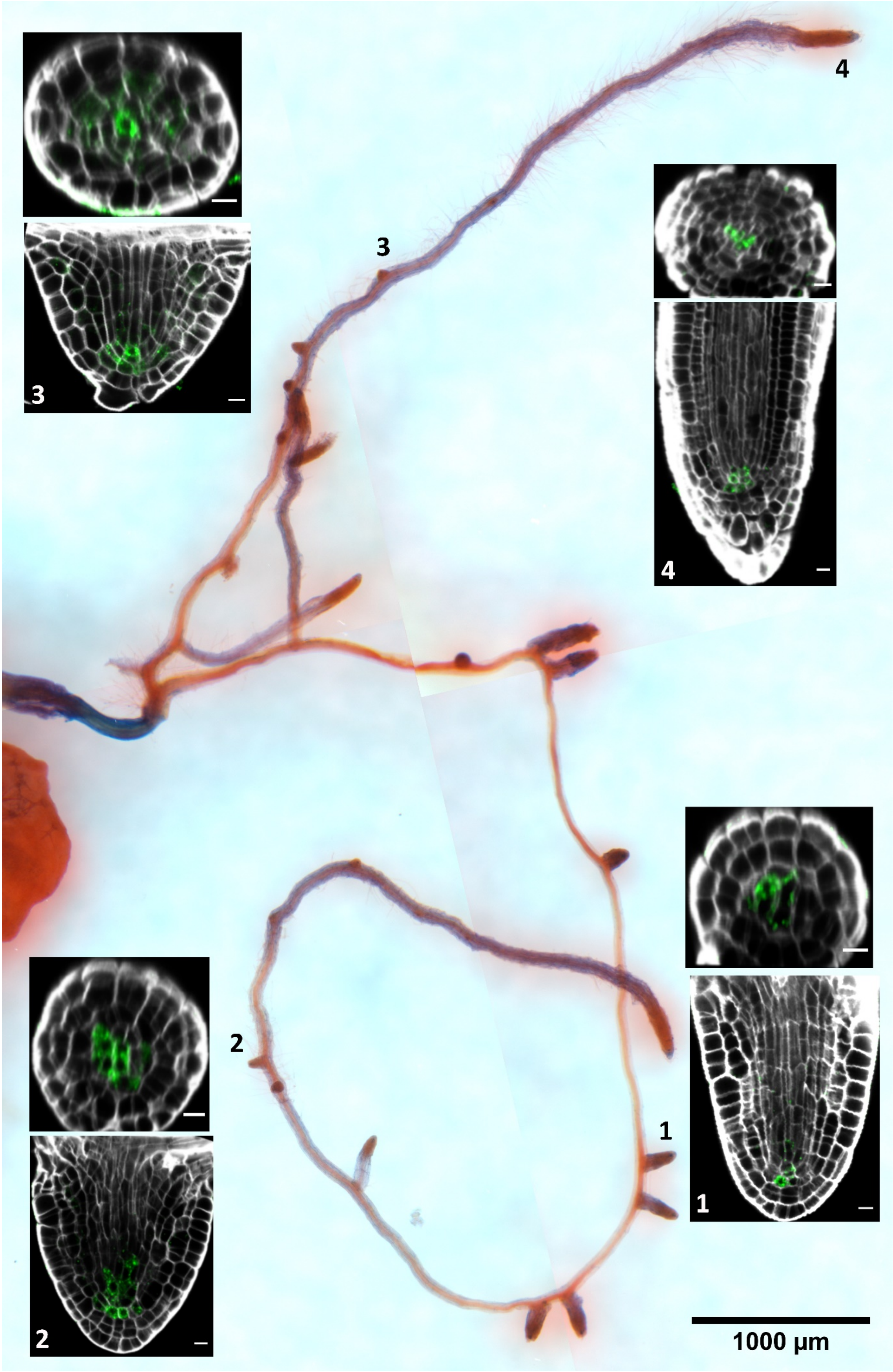


Figure S3

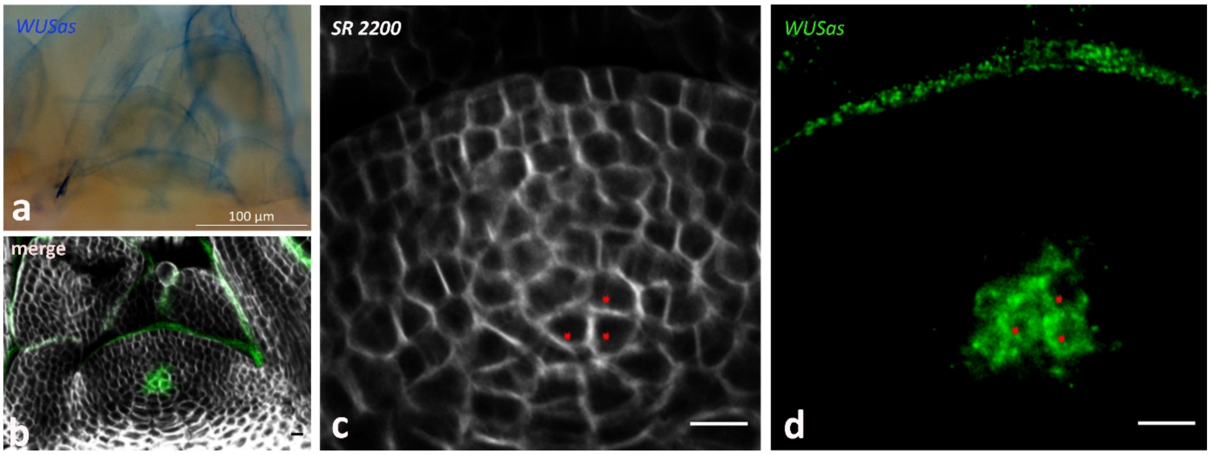


Figure S4

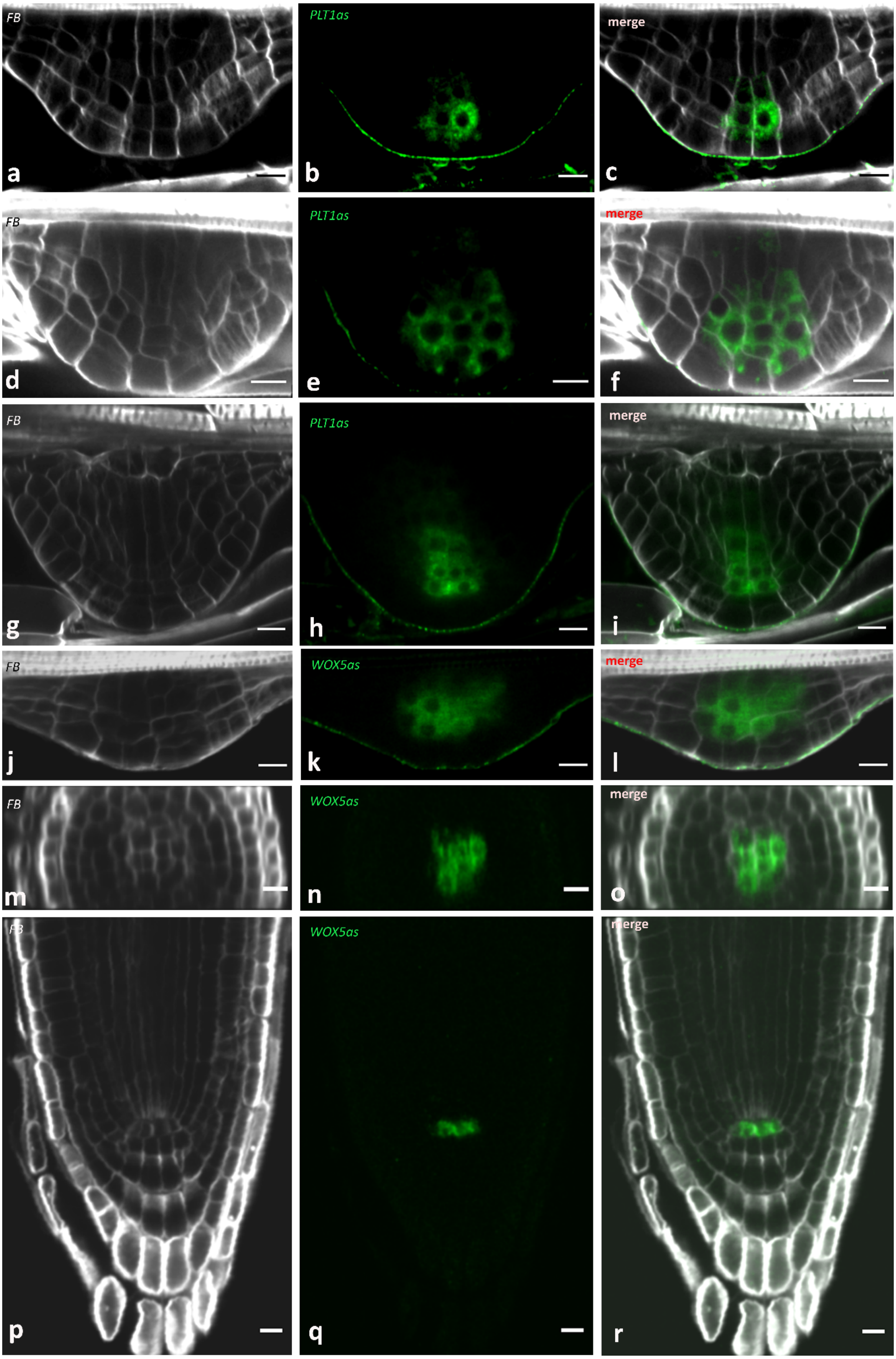
FB
a
PLT1as
b
merge
c
FB
d
PLT1as
e
merge
f
FB
g
PLT1as
h
merge
i
FB
j
WOX5as
k
merge
l
FB
m
WOX5as
n
merge
o
p
WOX5as
q
r

Figure S5

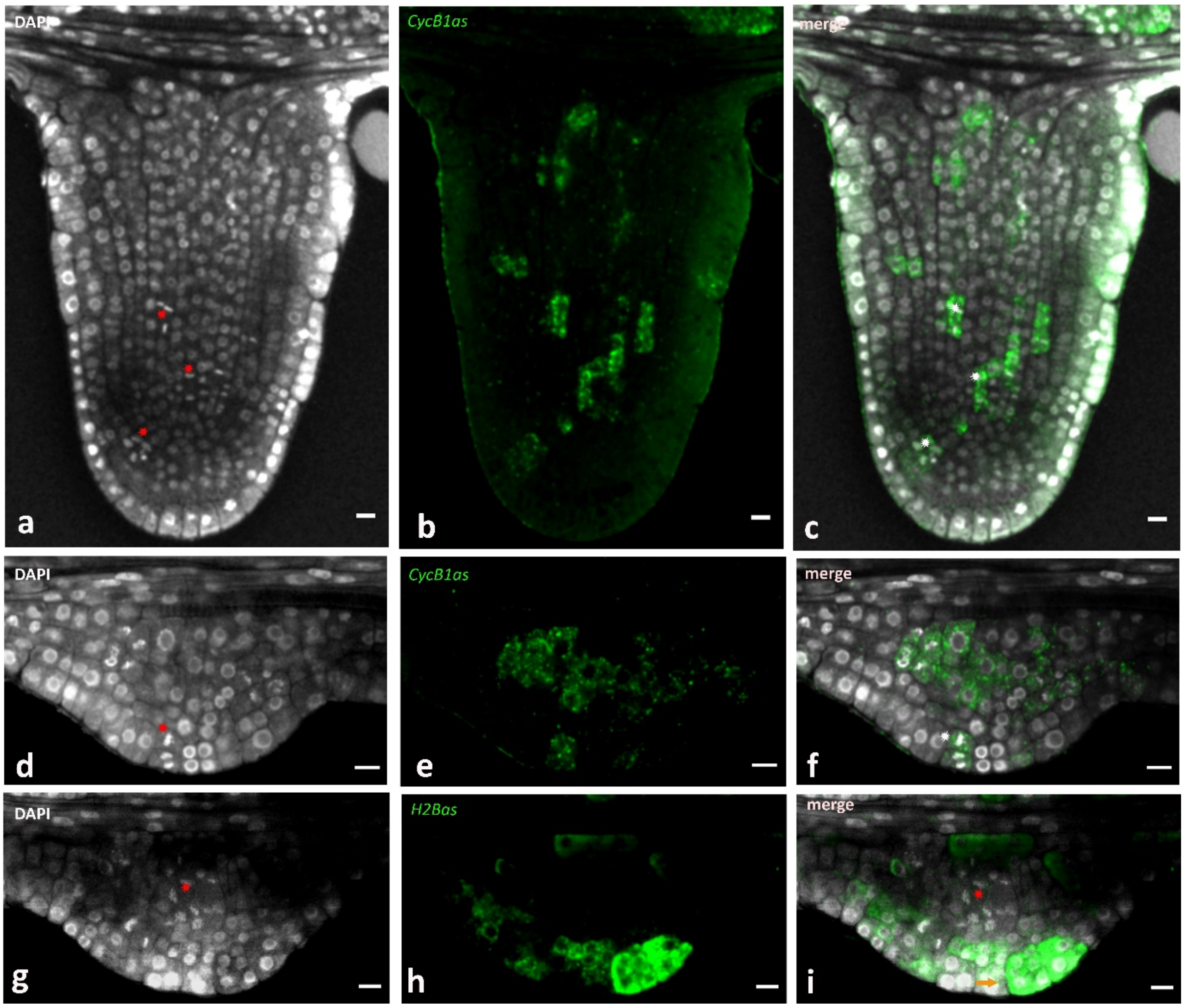
DAPI
CycB1as
merge
a
b
c
DAPI
CycB1as
merge
d
e
f
DAPI
H2Bas
merge
g
h
i

Figure S6

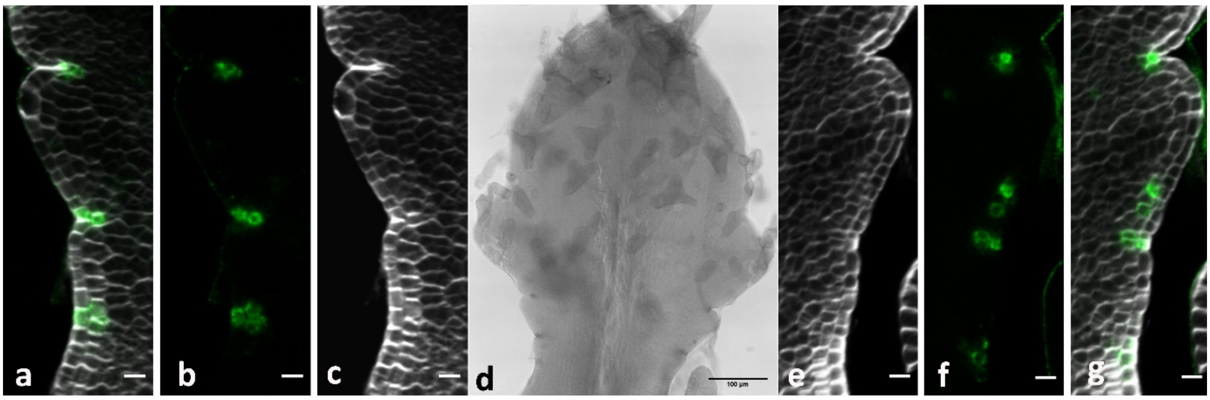


Figure S7

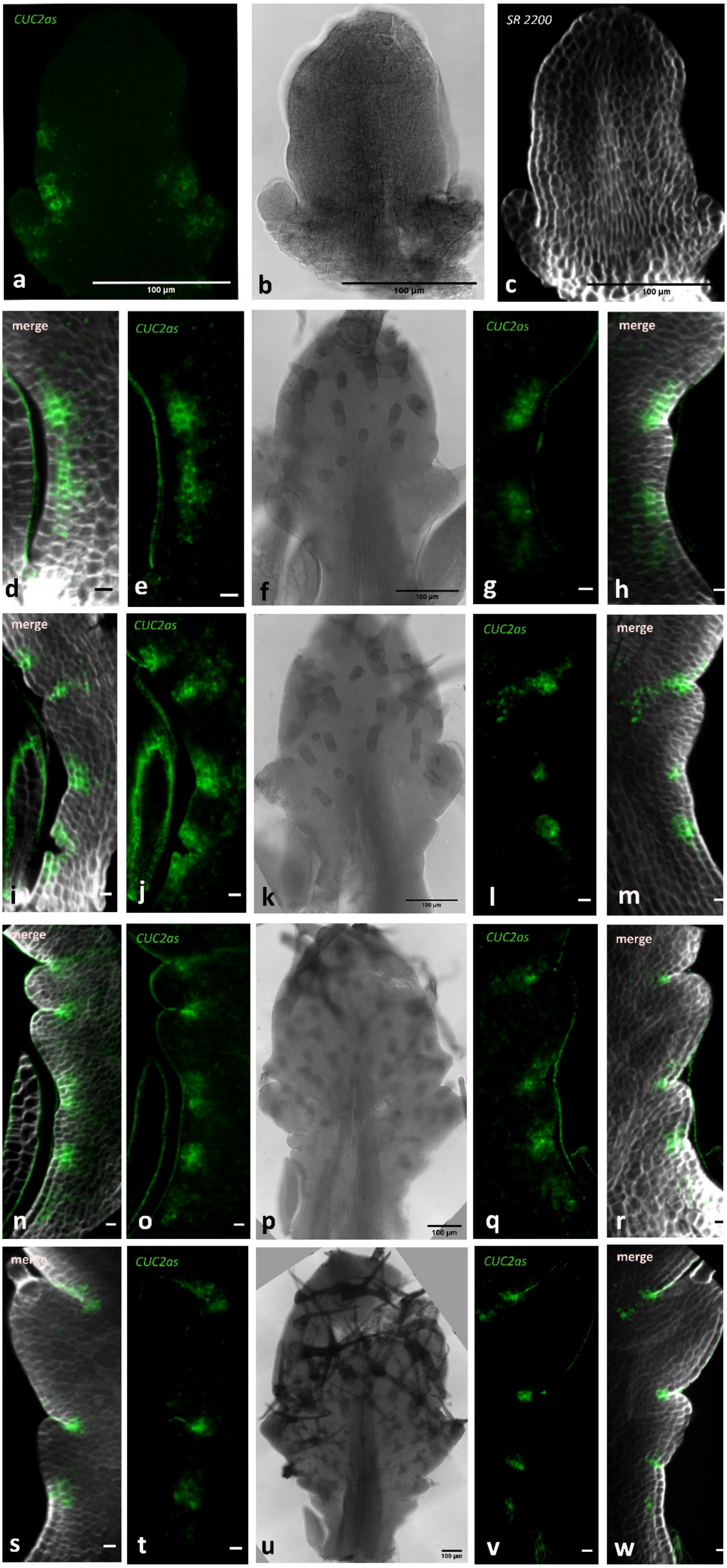

CUC2as
a
100 µm
b
100 µm
SR 2200
c
100 µm
merge
d
CUC2as
e
f
100 µm
CUC2as
g
merge
h
merge
i
CUC2as
j
k
100 µm
CUC2as
l
merge
m
merge
n
CUC2as
o
p
100 µm
CUC2as
q
merge
r
merge
s
CUC2as
t
u
100 µm
CUC2as
v
merge
w


Figure S8

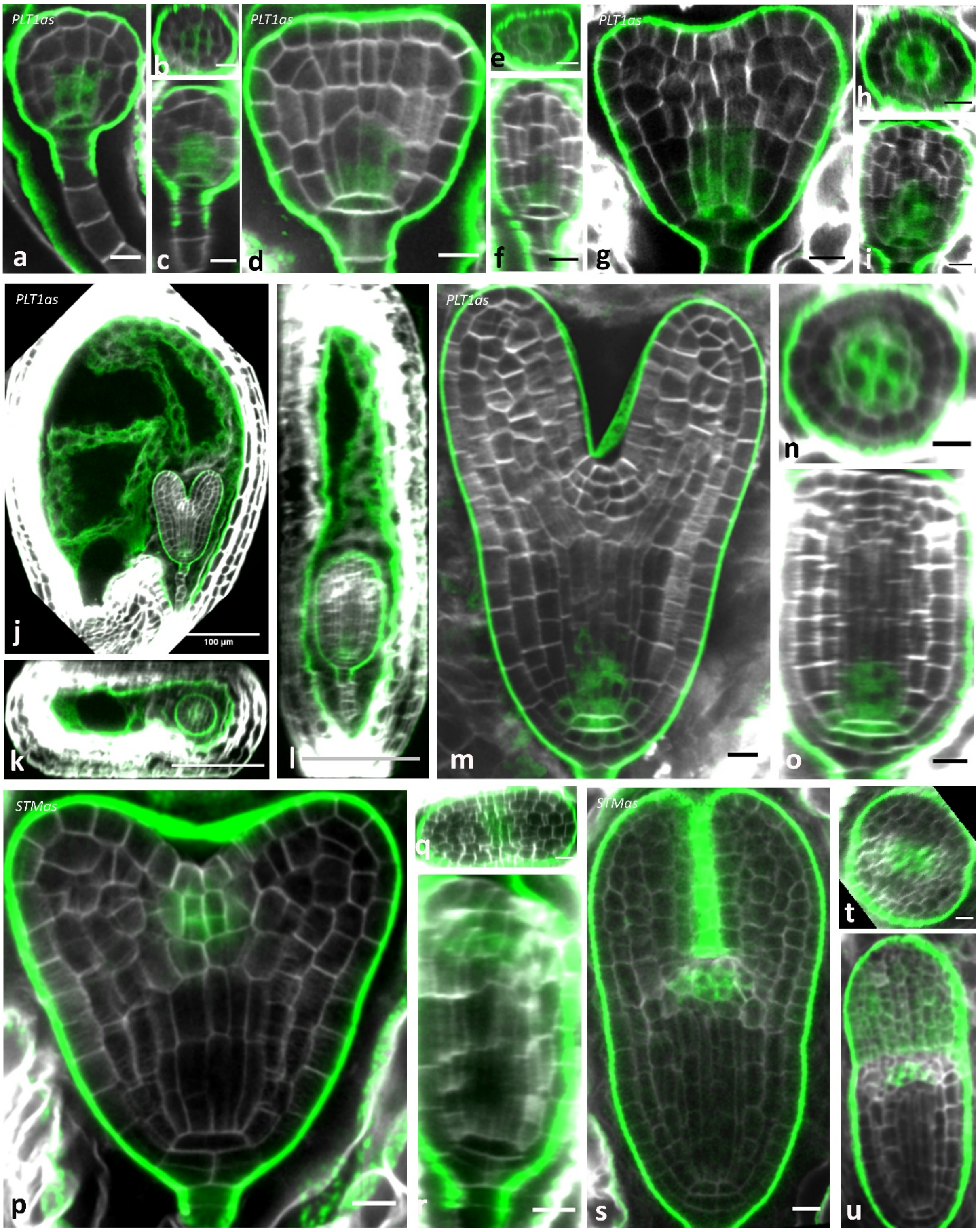


Figure S9

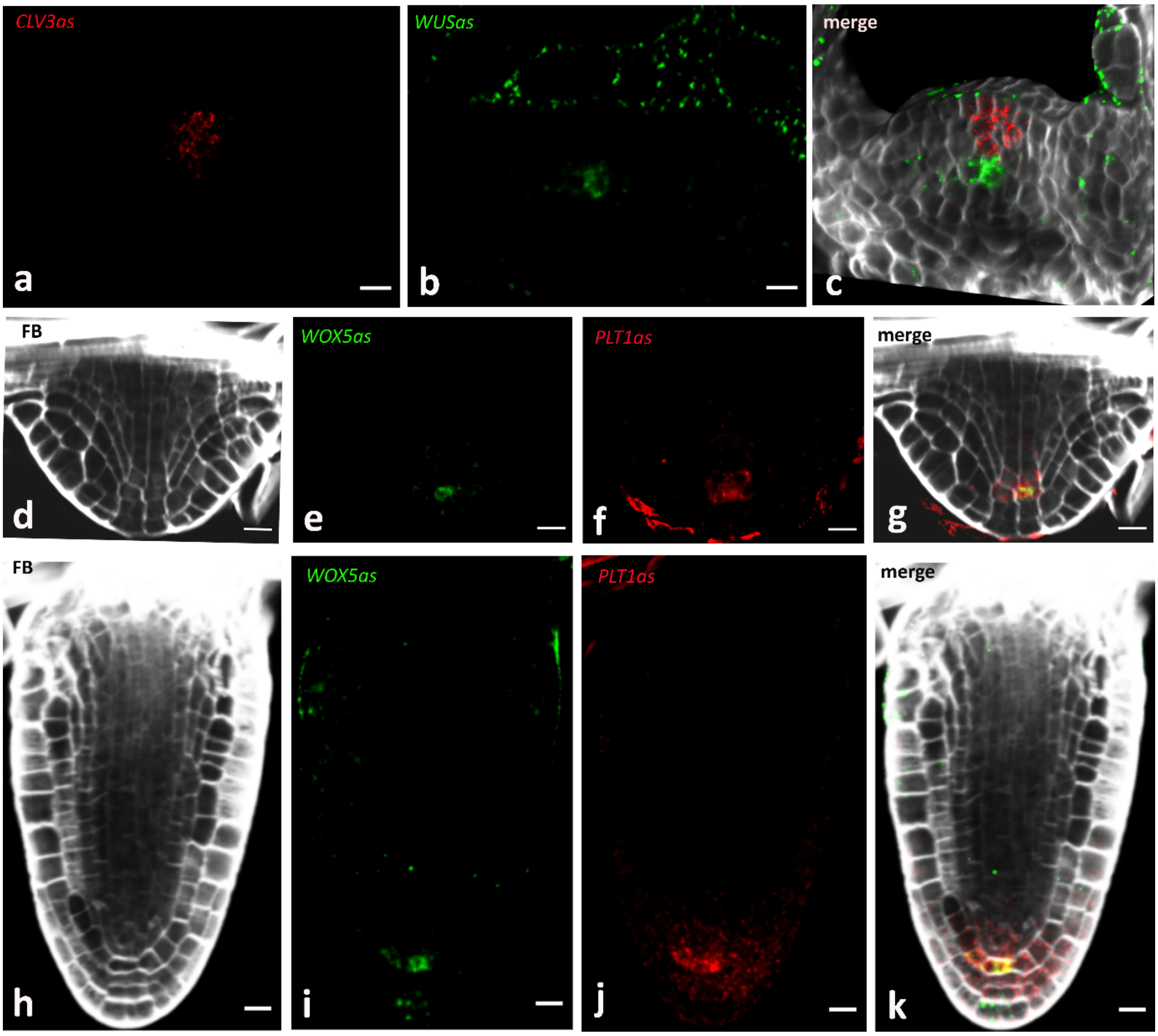

CLV3as
a
WUSas
b
merge
c
FB
d
WOX5as
e
PLT1as
f
merge
g
FB
h
WOX5as
i
PLT1as
j
merge
k


Figure S10

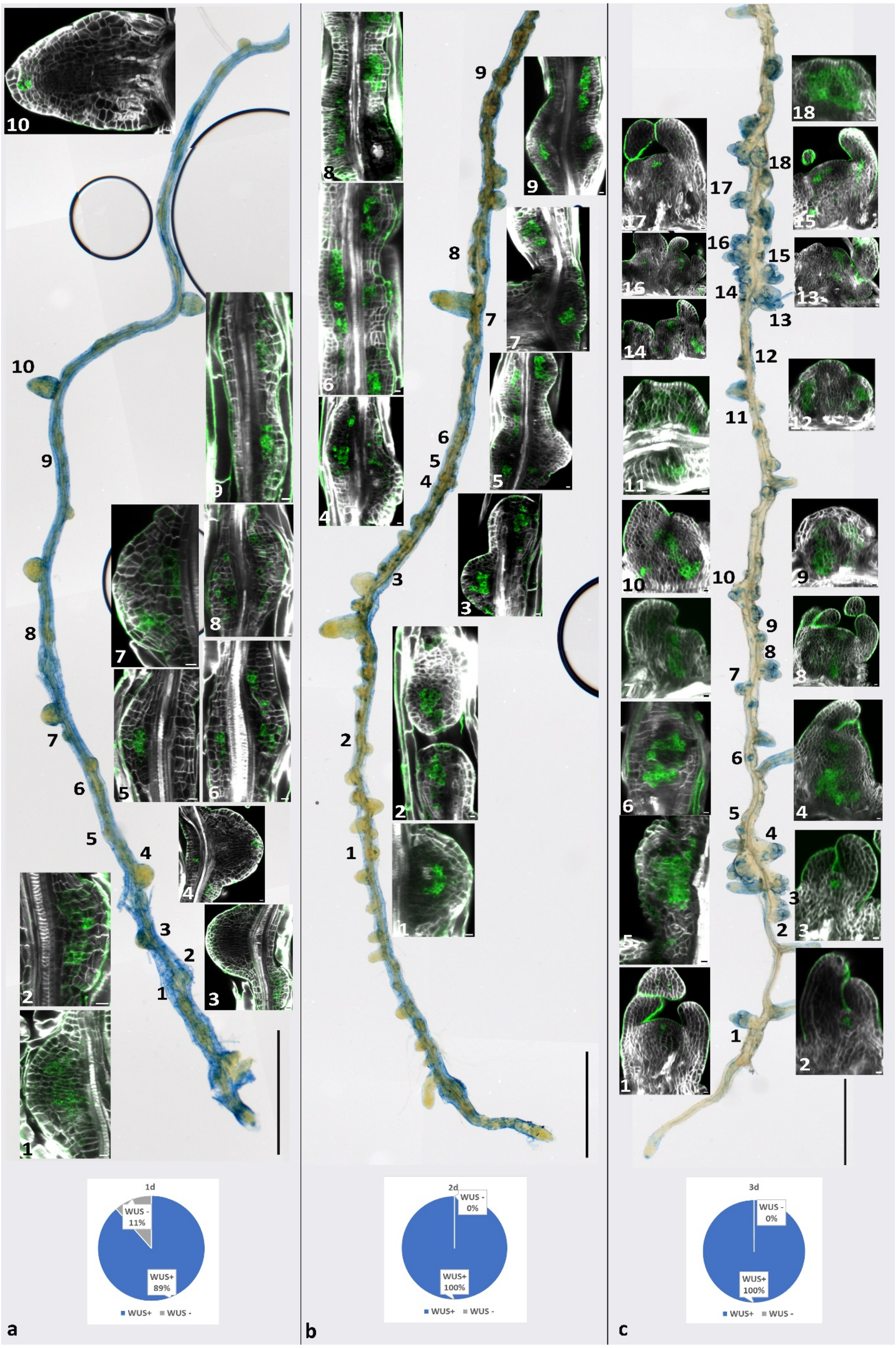


Figure S11

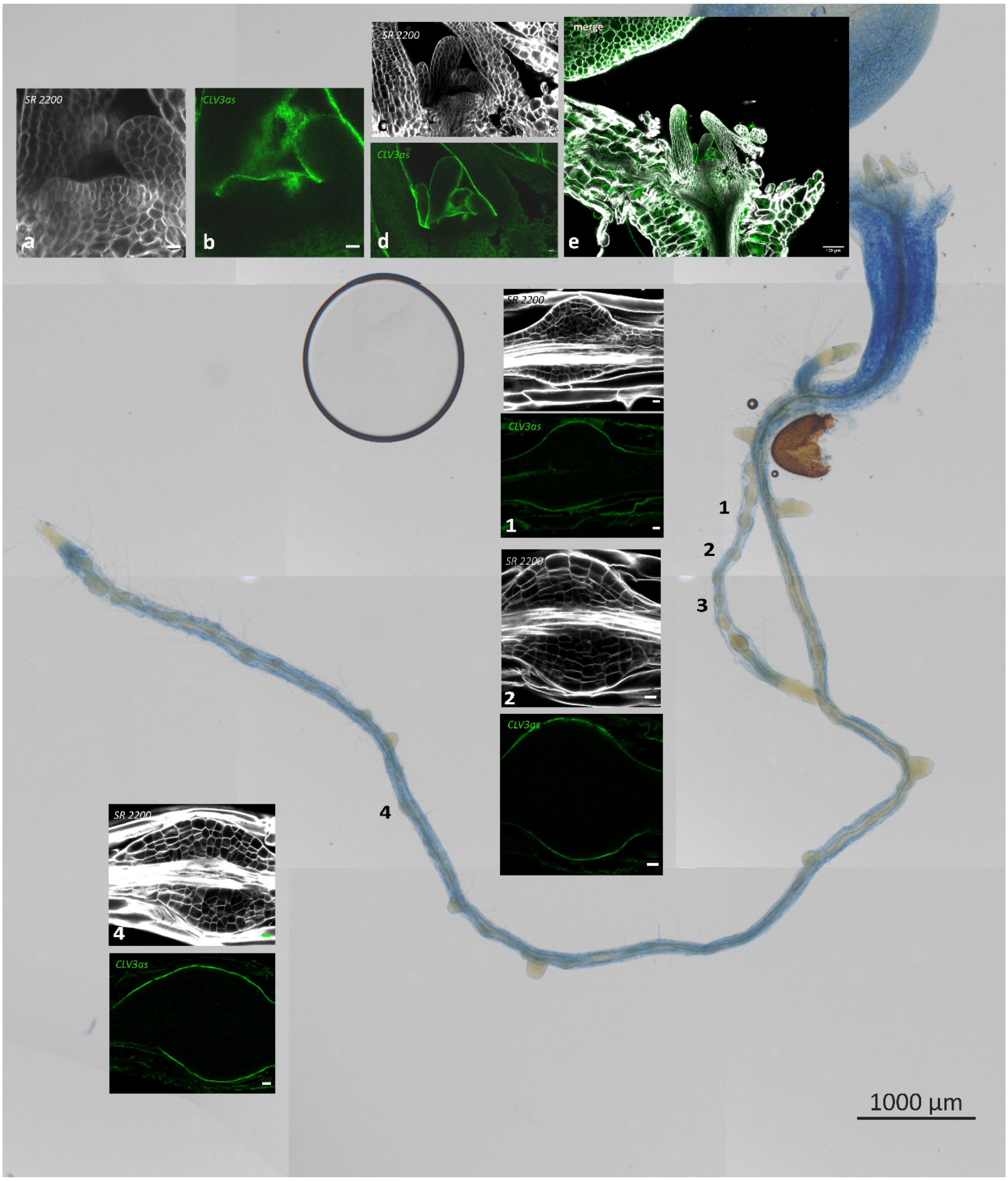


Figure S12

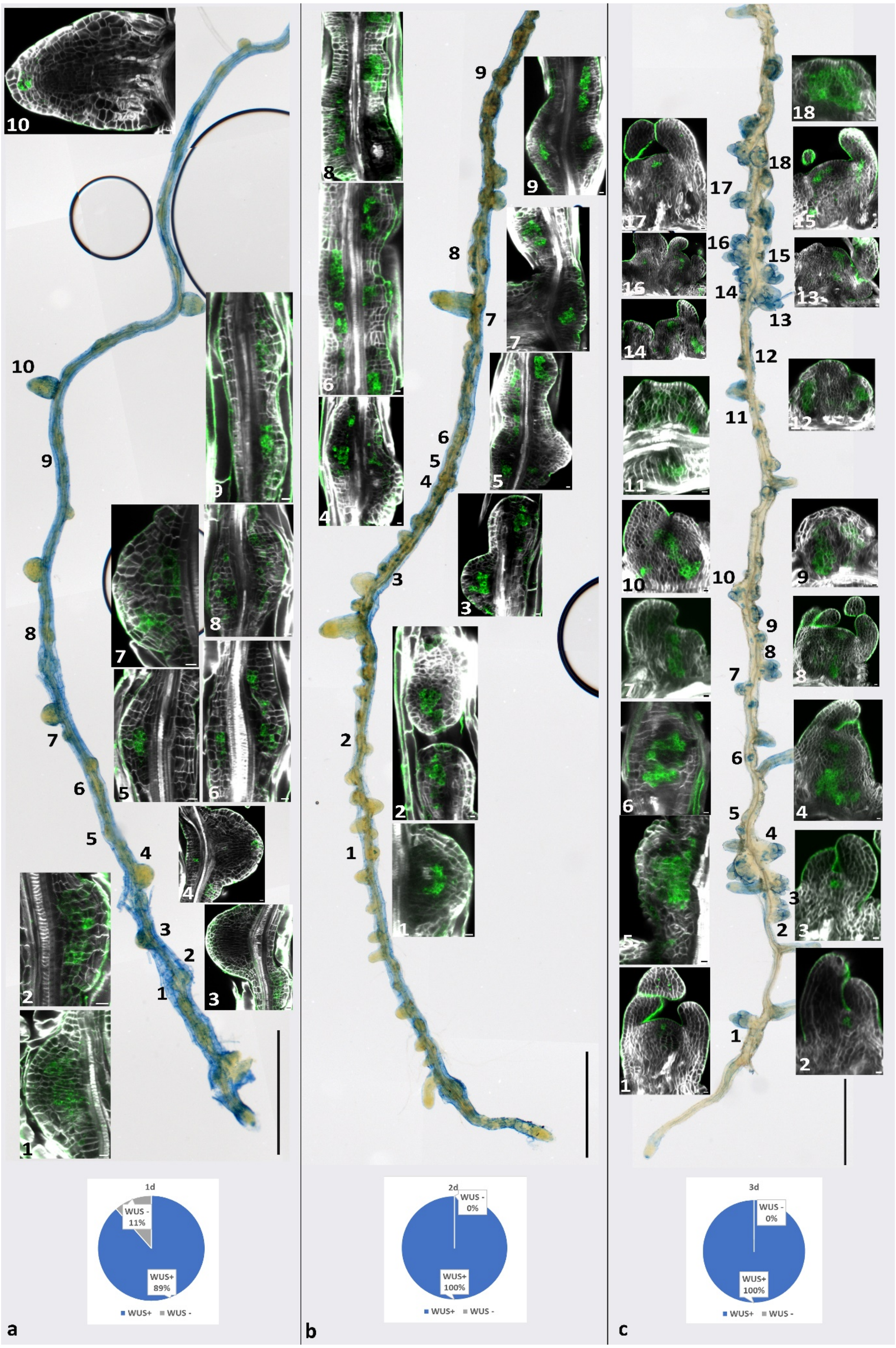
1d
WUS -
11%
WUS+
89%
WUS+ WUS -
2d
WUS -
0%
WUS+
100%
WUS+ WUS -
3d
WUS -
0%
WUS+
100%
WUS+ WUS -
a
b
c

Figure S13

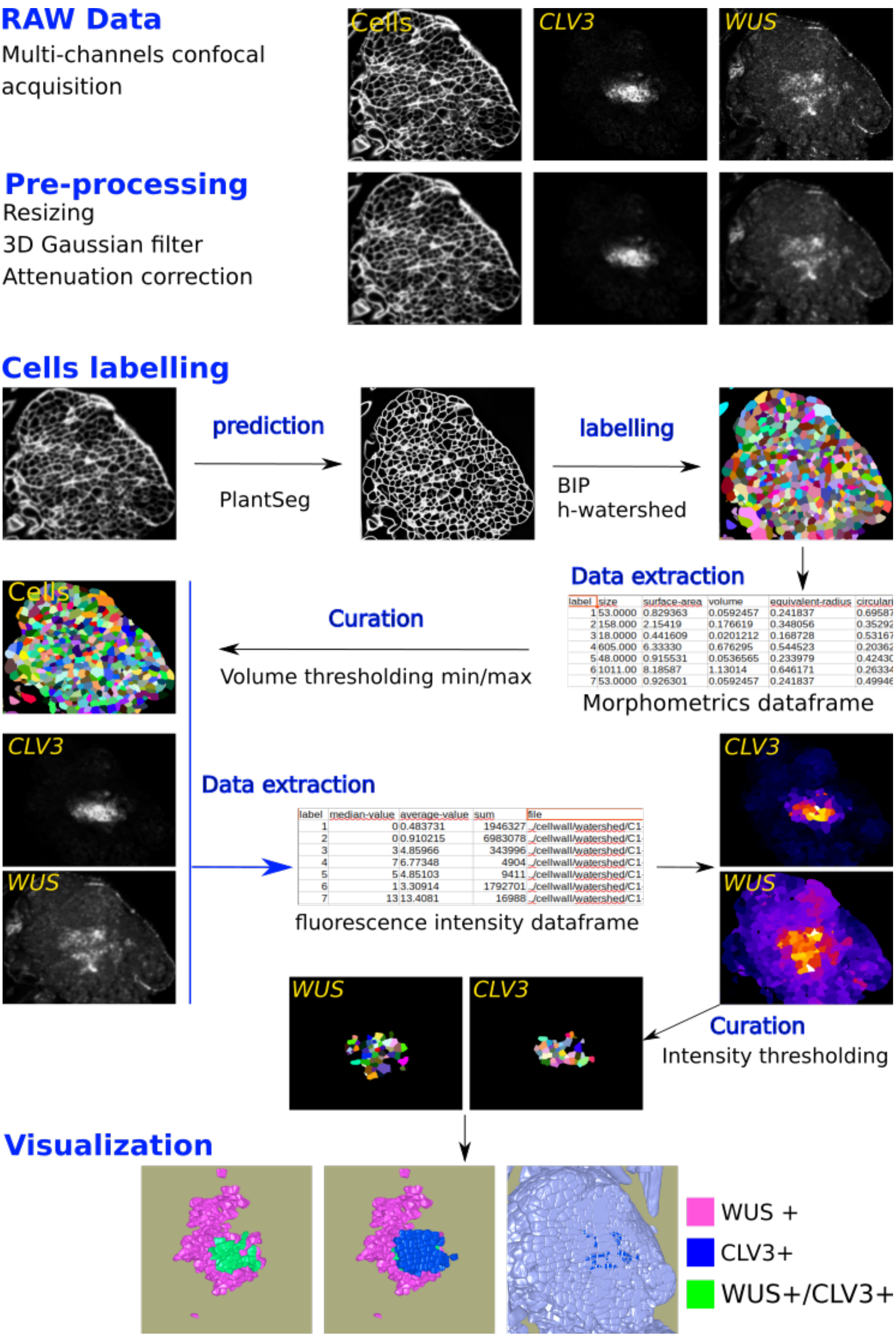
RAW Data
Multi-channels confocal acquisition
Cells
CLV3
WUS
Pre-processing
Resizing
3D Gaussian filter
Attenuation correction
Cells labelling
prediction
PlantSeg
labelling
BIP
h-watershed
Data extraction
Morphometrics dataframe
Curation
Volume thresholding min/max
Cells
CLV3
WUS
Data extraction
fluorescence intensity dataframe
CLV3
WUS
Curation
Intensity thresholding
WUS
CLV3
Visualization
WUS +
CLV3+
WUS+/CLV3+

Figure S14
