## Supplementary material for "M2WISH: an easy and efficient protocol for whole-mount mRNA *in situ* hybridization that allows 3D cell resolution of gene expression in *Arabidopsis thaliana*": detailed protocol

Reagents:

Digoxigenin-11-UTP (Roche, Cat No: 11209256910)

Fluorescein-12-UTP (Roche, Cat No: 11373242910

Ribonucleotide Triphosphates (rNTPs) (Promega, Cat No: P1221)

T7 RNA Polymerase (Promega, Cat No: P2075)

RNasin® Ribonuclease Inhibitors (Promega, Cat No: N2611)

Nuclease-Free Water (Promega, Cat No: P1193)

RQ1 RNase-Free DNase (Promega, Cat No: A1542)

Ribonucleic acid, transfer from baker's yeast (Roche, Cat No: 10109495001)

Ammonium acetate (Sigma-Aldrich, Cat No: P6148)

Sodium acetate (Sigma-Aldrich, Cat No: S2889)

Acetic acid, glacial (Sigma-Aldrich, Cat No: 695092)

Ethyl alcohol, Pure (Sigma-Aldrich, Cat No: 493511)

Paraformaldehyde (Sigma-Aldrich, Cat No: P6148)

Triton™ X-100 (Sigma-Aldrich, Cat No: T9284)

PBS 10X Buffer (Eurobio Scientific, Cat No: GAUPBS00-01)

Proteinase K, recombinant, PCR Grade (Roche, Cat No: 3115836001)

Glycine (Sigma-Aldrich, Cat No: G7126)

Triethanolamine (Sigma-Aldrich, Cat No: 90279)

Acetic anhydride (Sigma-Aldrich, Cat No: 320102)

Hydrochloric acid (Sigma-Aldrich, Cat No: H1758)

RCL2® (Exilone, Cat No: R1505J)

Formamid Deonized (Eurobio Scientific, cat No: GHYFORO1-01)

SSC 20X Buffer (Eurobio Scientific, Cat No: GHYSSC00-07)

TWEEN® 20 (Sigma-Aldrich, Cat No: 8.22184)

Denhardt's solution, 100x Concentrate (CliniSciences, Cat No: D0062)

Dextran sulphate sodium salt, M.W. ~ 500,000(VWR Chemicals, Cat No: 0198)

Sodium dodecyl sulfate solution (Eurobio Scientific, Cat No: GHYSDS02)

Ribonuclease A from bovine pancreas (Sigma, Cat No: R6513)

EDTA (Sigma-Aldrich, Cat No: E3889)

NaCl (Sigma-Aldrich, Cat No: S3014)

Macerozyme R10 (Duchefa, Cat No: M8002.0005)

Cellulose RS (Duchefa, Cat No: C8003.0005)

Pectolyase Y23 (Duchefa, Cat No: P9004.0001)

Pectinase (Serva, Cat No: 31660)

BSA FRACTION V (Eurobio Scientific, Cat No: GAUBSA01-62)

TRIS (Eurobio Scientific, Cat No: GAUTRI00-66)

Anti-Digoxigenin-AP, Fab fragments (Roche, Cat No: 11093274910)

Anti-Fluorescein-AP, Fab fragments (Roche, Cat No: 11426338910)

Anti-Digoxigenin-POD, Fab fragments (Roche, Cat No: 11207733910)

Vector® Blue Substrate Kit, Alkaline Phosphatase (AP) (Vector Laboratories, Cat No: SK-5300)

Vector® VIP Substrate Kit, Peroxidase (HRP) (Vector Laboratories, Cat No: SK-4600)

Invitrogen™ Alexa Fluor™ 488 Tyramide SuperBoost™ Kit, goat anti-mouse IgG (Fisher Scientific, Cat No: 15611892)

Sodium deoxycholate (Sigma-Aldrich, Cat No: D6750)

OmniPur® Urea (Sigma-Aldrich, Cat No: 9510-OP)

Xylitol (Sigma-Aldrich, Cat No: W507630)

Fluorescent Brightener 28 (Sigma-Aldrich, Cat No: F3543)

Direct Red 23 (Sigma-Aldrich, Cat No: 212490)

SR 2200 Cell Wall Stain (Renaissance Chemicals)

DAPI (Roche, Cat No: 10236276001)

Citifluor Antifadent Mountant Solutions (Agar Scientific, Cat No: AGR1320)

Primers for *in situ* hybridization

| | |
|---|---|
| CLV3-F | AAAAATGGATTCGAAGAGTTTTCTGC |
| CLV3-R-T7 | TGTAATACGACTCACTATAGGGCAAGAGATTAGGTCAAGGGAGCTGA |
| CUC2 -F | ATGGACATTCCGTATTACCA |
| CUC2-R-T7 | TGTAATACGACTCACTATAGGGCTCAGTAGTTCCAAATACA |
| CUC3-F | ATGATGCTTGCGGTGGAAGA |
| CUC3-R-T7 | TGTAATACGACTCACTATAGGGCCTACAGCTGGAATCCTAAA |
| CycB1-F | AGACGCCCCCACTACTTAGA |
| CycB1-R-T7 | TGTAATACGACTCACTATAGGGCTCGAGCAGCAACTAAACCAA |
| H2B-F | GCCGAGAGCAGAGAAGAAGC |
| H2B-R-T7 | TGTAATACGACTCACTATAGGGCTTGGTAACAGCCTTGGTTCC |
| PIN1-F | GGTCCTGGAGAAGCTGTGTTTGG |
| PIN1-R-T7 | TGTAATACGACTCACTATAGGGTGTTGGTGGCATCACCTTAGCC |
| PLT1-F | GCCGGAAACAAAGACCTCTA |
| PLT1-R-T7 | TGTAATACGACTCACTATAGGGCTGTTGGTCTGTTGGTGGAGA |
| STM-F | GGTTGTGGCGAGGCTAGAGG |
| STM-R-T7 | TGTAATACGACTCACTATAGGGCTCAAAGCATGGTGGAGGA |
| WOX5-F | TCTCCGTGAAAGGTCGAAGC |
| WOX5-R-T7 | TGTAATACGACTCACTATAGGGCATGGCGGTGGATGTTCCATT |
| WUS-F | GGCTGAGACAGTTCGGAAAG |
| WUS-R-T7 | TGTAATACGACTCACTATAGGGCCCAAACAGAGGCTTTGCTCT |
| PLT1-F-T7 | TGTAATACGACTCACTATAGGGCTGTTGGTCTGTTGGTGGAGA |
| PLT1-R | GCCGGAAACAAAGACCTCTA |

Fixative Solution:

1. Add paraformaldehyde powder to 1xPBS to final concentration 4%.
2. Stir the solution in 80°C water bath until a complete dissolving.
3. Makes 10 ml aliquots and freeze. Aliquots can be stored at -20°C for years.
4. Thaw PFA aliquot, add 100µl of 10% Triton X100 and 10µl of SR2200. This aliquot can't be reused.

Hybridization solution:

For 10 ml of hybridization solution mix the following:

| | |
|---|---|
| Formamide | 5 ml |
| 50% Dextran sulfate | 2 ml |
| tRNA (10mg/ml) | 1 ml |
| Tween 20 | 750 µl |
| 100x Denhardt | 250 µl |
| Sterile water | 1 ml |

NOTE: Hybridization solution should be prepared in advance as mixing produces a lot of bubbles.

ClearSee solution is prepared according to (Tofanelli *et al.*, 2019):

- Xylitol [final 10% (w/v)]

- Sodium Deoxycholate [final 15% (w/v)]

- Urea [final 25% (w/v)]

- Water to the final volume

NOTE: Mix the ClearSee solution on the magnetic stirrer with heating until everything is completely dissolved.

### WISH Protocol

#### DAY1

1. Fixe samples under vacuum for 1 hour at room temperature (RT), then over/night (O/N) at 4°C. For cell wall staining, SR2200 can be added direct to fixative at final concentration 0.1%. Do not leave samples more than 2 days in fixative.

#### DAY2

2. Permeabilize samples by microwave treatment in series of Ethanol/water dilutions: 10%, 30%, 50%, 70%. Apply 180W 6 times for 30s (or up to boiling) in each bath. After this step, samples can be stored in 70% ethanol for years at -20°C.
3. Rehydrate in Ethanol/water dilution: 70%, 50%, 30%, 10%, then PBS-Tween 0.1% 5min each.
4. Digest with proteinase K for 10min at RT (use final concentration 10µg.ml $^{-1}$ in PBS).
5. Stop the digest with glycine (2mg.ml$^{-1}$) in PBS for 2min at RT.
6. Wash samples in Triethanolamine/ Acetic anhydride 10min at RT with a gentle shaking. For working solution, dilute 300µl of Triethanolamine in 20ml of sterile water, add 70µl of 37% HCl, then add 1ml of Acetic anhydride. The reaction starts when Acetic anhydride is added.
7. Fix samples in RCL2 for 15min in vacuum.
8. Wash three times in PBS-Tween for 1min at RT.
9. Distribute samples in baskets in a 24-well plate containing PBS-Triton.
10. Prehybridize samples in 50% formamide, 5xSSC, 0.1% Tween-20, for 60min at 47°C in a water bath. Add 500µl of pre-hybridization solution per well.
11. **For one probe detection:**

    Dilute RNA probe by halving with DEPC treated water (Choose the volume of probe according to labelling efficiency*). Denature RNA probe for 2min at 80°C, chill on ice and mix with 400µl of hybridization solution per well.

    **For two probes detection:**

Dilute both RNA probes by halving with DEPC treated water (Choose the volume of probe according to labelling efficiency*). Denature RNA probes for 2min at 80°C, chill on ice and mix them with 400µl of hybridization solution per well.

12. Hybridize at 47°C O/N.

#### DAY3

13. Wash in 0.1xSSC, 0.5% SDS for 30min at 47°C.
14. Wash in 2xSSC, 50% Formamide for 60min at 47°C.
15. Wash in TNE (Tris 10mM, EDTA 1mM, NaCl 100mM, pH 8.0) buffer for 5min at 47°C.
16. Incubate with RNAse A for 30min at 37°C (use final concentration 20µg.ml$^{-1}$ in TNE).
17. Wash in TNE for 5min at 47°C.
18. Wash in 2xSSC, 50% Formamide for 60min at 47°C.
19. Wash in 0.1xSSC for 2min at 47°C.
20. Leave samples in PBS for O/N at RT or proceed directly to step 21.

#### DAY4

21. Perform a partial cell wall digestion with an enzymatic mix (0.08%macerozyme; 0.08% cellulase; 0.04% pectolyase; 0.12% pectinase) for 10min at RT.
22. Wash in PBS three times for 5min at RT.
23. Pre-incubate in 10mM Tris-HCl pH7.5; 15mM NaCl, 1% BSA, 0.5% Triton X-100 for 60min at RT.

    **For one probe detection:**

24. Incubate samples with Anti-Dig-Fab-AP (1/1250 in a above solution O/N at 4°C or 3 hours at 37°C.
25. Wash in 100mM Tris-HCl pH8.2; 0,1% Tween-20 three times for 10min at RT.
26. Perform AP staining reaction with Vector® Blue Substrate Kit (Vector laboratories SK-5300 3x10min RT) according to manufactured protocol:

    for 5ml of 100mM Tris-HCl pH8.2; 0,1% Tween-20 add
    80µl of Reagent 1
    80µl of Reagent 2
    45µl of Reagent 3
    Mix well before use, incubate with a gentle agitation in a dark at RT.
    The time of staining can last up to 2 days.

27. Incubate samples in ClearSee (Xylitol 10%; Deoxycholic acid 15%; Urea 25%) for at least 5 days, one of cell wall dye can be added to ClearSee to final concentration: SR2200 1/1000, 50µg.ml$^{-1}$ Fluorescent brightening, 0.01% Direct Red 80.

    Samples can be stored in Clearsee for months at 4°C.

28. Mount samples in ClearSee and proceed with confocal microscope analysis. Vector® Blue Substrate excitation was done at 633nm; its emission was collected at 740 to 800nm.

    SR2200 was excited with a 405nm UV laser and its emission recorded between 420 and 480nm

    **For two probes detection:**

29. Incubate samples with Anti-Dig-Fab-AP and Anti-Fluorescein-Fab-POD each in 1/1250 dilution O/N at 4°C or 3 hours at 37°C.

30. Wash three times in PBS-Triton 0.1% for 15min at RT.

31. Perform the fluorescent POD staining reaction with Tyramide Signal Amplification Kit according to (Rozier *et al.*, 2014):

    Prepare solution 1 by adding 1µl of $H_2O_2$ to 200µl of amplification buffer,

    Prepare solution 2 by adding 1µl of solution 1 to 100µl of amplification buffer.

    Prepare solution 3 by adding 12µl of Alexa Fluor 488 tyramide to 400µl of solution2.

    Transfer the samples to solution 3 and incubate them in the dark for 30min at RT with gentle agitation.

32. Wash three times in 100mM Tris-HCl pH8.2; 0,1% Tween-20for 10min at RT.

33. Perform AP staining reaction with Vector® Blue Substrate Kit (Vector laboratories SK-5300) according to manufactured protocol:

    for 5ml of 100mM Tris-HCl pH8,2; 0,1% Tween-20 add
    80µl of Reagent 1
    80µl of Reagent 2
    45µl of Reagent 3
    Mix well before use, incubate with a gentle agitation in a dark at RT.
    The time of staining can last up to 2 days.

34. Wash three times in 100mM Tris-HCl pH8.2; 0,1% Tween-20 for 10min at RT.

35. Keep samples in PBS. ClearSee solution is not appropriate for Alexa fluorochromes.

36. Add one of cell wall dyes to samples in PBS (Fluorescent Brightening to 50 µg.mL$^{-1}$ dilution, Direct Red 80 to 0.01% dilution, SR2200 to 1/1000 dilution). Incubate O/N at RT with a gentle shaking.

**37.** Mount samples in Citifluor AF1/PBS solution (3:1) with (optional a DAPI to final concentration 10μg.ml$^{-1}$) and proceed with confocal microscope analysis:
SR2200 excitation is done at 405 nm and its emission is detected at 420 - 480nm.
Alexa Fluor 488 excitation is done 488 nm and its emission is detected at 500 to 550nm.
Vector® Blue Substrate excitation is done at 633nm, its emission is detected between 740 to 800nm.

**Cell Wall Staining protocols compatible with WISH**

**1st option:** add Renaissance staining SR2200 to fixative (**step 1**) to 1/1000 dilution.

**2nd option**: add one of cell wall dyes to samples in Clearsee (Fluorescent Brightening to 50 μg.ml$^{-1}$ dilution, Direct Red 80 to 0.01% dilution, SR2200 to 1/1000 dilution). Incubate O/N at RT.

**3rdoption:** add one of cell wall dyes to samples in PBS (Fluorescent Brightening to 50μg.ml$^{-1}$ dilution, Direct Red 80 to 0.01% dilution, SR2200 to 1/1000 dilution). Incubate O/N at RT with a gentle shaking.

**Nuclei staining protocol compatible with WISH**

Wash samples three times in PBS for 5min.

Mount samples in Citifluor AF1/PBS solution (3:1) with a DAPI to final concentration 10μg.ml$^{-1}$.

**Probe labeling by *in vitro* transcription from a PCR amplified fragment:**

1. Prepare a PCR template using 3' primers extended with a sequence of T7 promoter.
2. Set up a reaction for *in vitro* transcription as tabulated below:

| | |
|---|---|
| DNA template (1μg) up to | 9μl |
| 5xBuffer | 4μl |
| ATP | 1.5μl |
| GTP | 1.5μl |
| CTP | 1.5μl |
| UTP | 0.9μl |
| DIG (F)-UTP | 0.6μl |
| RNAse inhibiter | 1μl |
| 7RNA polymerase | 2μl |
| Nuclease free water | Up to 20μl |

3. Incubate at 37°C for 1h30.
4. Add to reaction 2μl of RNAse free DNAse and 68μl of Nuclease free water.

5. Incubate for 30min at 37°C.
6. Precipitate probe as tabulated below:

| | | |
|---|---|---|
| tRNA (10mg/mL) | 10 µL | |
| 10M Ammonium Acetate | 20 µL | |
| EtOH absolute | 240 µL | O/N at -20°C |
| Centrifuge | 13000 rcf | 30 min 4°C |
| Wash pellet with EtOH 70% | 13000 rcf | 10 min 4°C |

7. Dry pellet.
8. Hydrolyze pellet in 50µL of carbonate buffer (120mM $Na_2CO$; 80mM $NaHCO_3$; pH 10.2) for 10min at 60°C, thus probes of different length are hydrolysed into 100 – 200 bp fragment.
9. Stop reaction by adding:

| | | |
|---|---|---|
| 10% acetic acid | 10 µL | |
| 3 M Sodium Acetate pH 4.8 | 12 µL | |
| 100% EtOH | 312 µL | O/N at -20°C |
| Centrifuge | 13000 rcf | 30 min at 4°C |
| Wash pellet with EtOH 70% | 13000rcf | 10 min at 4°C |

10. Dry and then dissolve pellet in 20µl of Nuclease free water and 20µl Formamide**.**

**Dot-Blot control of labeling efficiency**:

DIG or FITC sole detection

1. Prepare dilution series (1:10, 1:100, 1:1000) of the labeled probes applied 1µl of each dilution to a small strip of positively charged nylon membrane.
2. Preincubate membrane in 10mM Tris-HCl pH7.5; 15mM NaCl, 1% BSA, 0.5% Triton X-100 for 30min at RT.
3. Incubate membrane with Anti-Digoxigenin-AP, Fab fragments (or Anti-FITC-Fab-AP) diluted 1/1250 in a above solution for 15min at RT.
4. Wash membrane three times in 100mM Tris-HCl pH8.2; 0.1% Tween-20 for 5min.
5. Perform AP staining reaction with Vector® Blue Substrate Kit (Vector laboratories SK-5300 3x10min RT) according to manufactured protocol (about 10min).
6. Wash in deionized water.

DIG and FITC simultaneous detection

1. Prepare dilutions series of both probes and applied 1µl of each dilution to the same positively charged nylon membrane.
2. Preincubate membrane in 10 mM Tris-HCl pH7.5; 15mM NaCl, 1% BSA, 0.5% Triton X-100 for 30min at RT.
7. Incubate membrane with Anti-Dig-Fab-AP and Anti-FITC-Fab-POD diluted 1/1250 in a above solution for 15min at RT.
3. Wash membrane three times in PBS-Triton 0.1% for 10min.
4. Perform POD staining reaction with Vector® VIP Peroxidase substrate kit according to manufactured protocol (about 5min).
5. Wash membrane three times in 100mM Tris-HCl pH8.2; 0.1% Tween-20 for 5min.
6. Perform AP staining reaction with Vector® Blue Substrate Kit (Vector laboratories SK-5300 3x10min RT) according to manufactured protocol (about 10min).
7. Wash in deionized water.
